# A unified framework for inferring the multi-scale organization of chromatin domains from Hi-C

**DOI:** 10.1101/530519

**Authors:** Ji Hyun Bak, Min Hyeok Kim, Lei Liu, Changbong Hyeon

## Abstract

Identifying chromatin domains (CDs) from high-throughput chromosome conformation capture (Hi-C) data is currently a central problem in genome research. Here we present a unified algorithm, Multi-CD, which infers CDs at various genomic scales by leveraging the information from Hi-C. By integrating a model of the chromosome from polymer physics, statistical physics-based clustering analysis, and Bayesian inference, Multi-CD identifies the CDs that best represent the global pattern of correlation manifested in Hi-C. The multi-scale intra-chromosomal structures compared across different cell types allow us to glean the principles of chromatin organization: (i) Sub-TADs, TADs, and meta-TADs constitute a robust hierarchical structure. (ii) The assemblies of compartments and TAD-based domains are governed by different organizational principles. (iii) Sub-TADs are the common building blocks of chromosome architecture. CDs obtained from Multi-CD applied to Hi-C data enable a quantitative and comparative analysis of chromosome organization in different cell types, providing glimpses into structure-function relationship in genome.

---

Chromosome conformation capture (3C) and its derivatives, which are used to identify chromatin contacts through the proximity ligation techniques (1, 2), take center stage in chromosome research (3, 4). Square-block and checkerboard patterns manifested in Hi-C data provide glimpses into the organization of chromatin chains inside cell nuclei. Despite cell-to-cell variations inherent to Hi-C data, which is in practice collected from a heterogeneous cell population, the cell-type specificity of chromosome architecture gleaned from Hi-C is still clear. Furthermore, the change of Hi-C pattern with the transcription activity and the phase of cell cycle underscores the functional roles of chromosome structure in gene regulation (5–14). Given that pathological states of chromatin are also manifested in Hi-C (15, 16), accurate characterization of chromatin domains (CDs) from Hi-C data is of great importance in advancing our quantitative understanding of the genome function.

Inside cell nuclei each chromosome made of 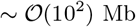 DNA is segregated into its own territory (Fig. 1a) (17). At the scale of 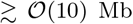, alternating blocks of active and inactive chromatin are phase-separated into two megabase sized aggregates, called A- and B-compartments (18–21) (Fig. 1b). Topologically associated domains (TADs), detected at 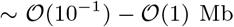 (22–25), are considered the basic functional unit of chromatin organization and gene regulation because of their well-conserved domain boundaries across cell/tissue types (17, 20, 26, 27). It was suggested that the proximal TADs in genomic neighborhood aggregate into a higher-order structural domain termed “meta-TAD” (6). At smaller genomic scale, each TAD is further split into sub-TADs that display more localized contacts (19, 28–31) (Fig. 1c).

**Fig. 1.**
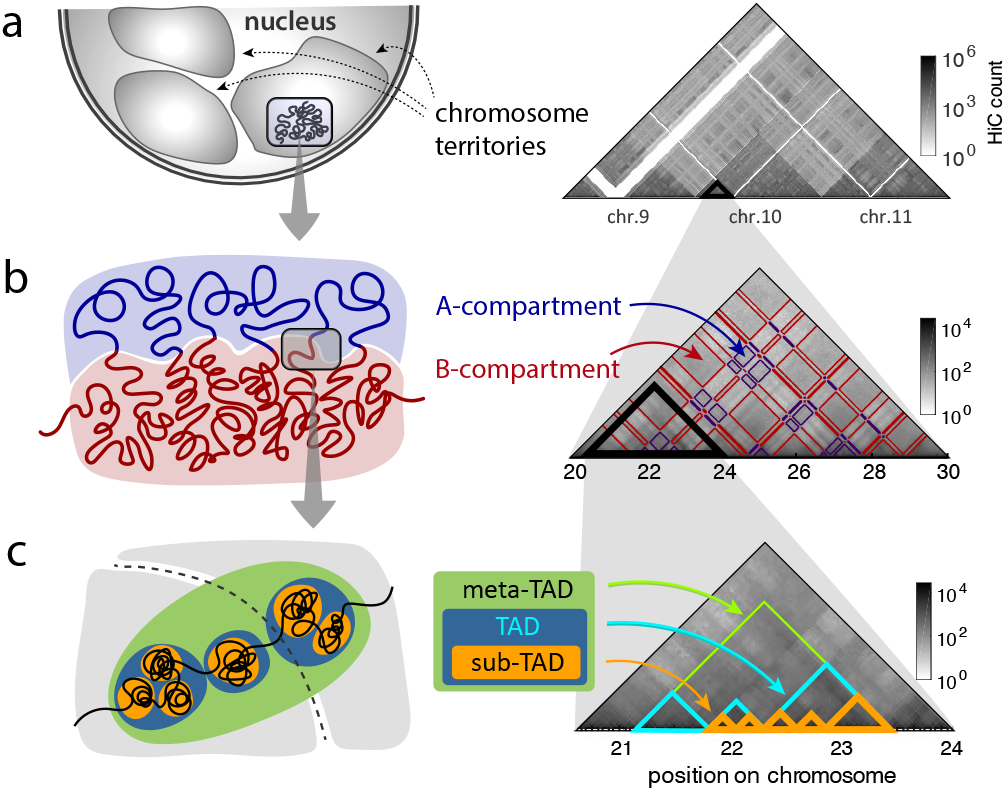
The hierarchical organization of interphase chromosome and Hi-C map. **(a)** Chromosome territories in the cell nucleus, which are manifested as the higher intra-chromosomal counts in the Hi-C map. **(b)** Alternating blocks of active and inactive chromatins, segregated into A- and B-compartments, give rise to the checkerboard pattern on Hi-C. **(c)** Sub-megabase to megabase sized chromatin folds into TADs. Adjacent TADs are merged to meta-TAD (6), and individual TAD is further decomposed into sub-TADs (19, 28–31).

Currently there are many algorithms available to identify CDs from Hi-C data, making important contributions to understanding the intra-chromosome architecture. However, CDs identified using different algorithms or parameters display significant variations, and there is no generally accepted definition for the above-mentioned CD at each scale. For example, the average size of a TAD varies from 100 kb to 2 Mb depending on the specific algorithm being used. Furthermore, still lacking is a unified algorithm to characterize CDs at multiple scales. In many of the existing algorithms specialized in finding CDs at particular genomic scales, Hi-C data should first be formatted at specific resolution (18, 19, 22). Although there are methods (32, 33) developed for identifying hierarchical domain structures of chromosomes, their algorithms rely on local pattern recognition analyses (18, 19, 22, 34), not implementing the physical viewpoint that chromosomes are a three dimensional object made of a long polymer (6, 14, 35–40).

Here, we interpret Hi-C data as pairwise contact probability matrix resulting from polymer networks whose inter-monomer distances are harmonically restrained. The cross-correlation matrix derived from Hi-C is used as the sole input for Multi-CD, the algorithm that we have developed to identify CD at varying genomic scales. Agreement of CD solutions from Multi-CD with the previous knowledge on chromatin organization as well as with information from bio-markers indicates the reliability of Multi-CD. A single algorithm-based solutions of CD allow us to assess the multi-scale structure of chromatin from a more unified perspective. We assert that amid the rapidly expanding volume of Hi-C data (10–12), Multi-CD holds good promise to quantitative and principled determination of chromatin organization.

## Theory

### Transforming Hi-C into a cross-correlation matrix using a polymer network model

A chromosome can be viewed as a polymer chain that is folded to a network structure characterized with multiple cross-links (35, 41, 42). Even in the interphase that displays less amount of activity than mitotic phase, continuous events of free energy consumption break the detailed balance condition, driving the chromosome out of equilibrium (43–47). However, chromosome dynamics in each phase is slow (39, 40, 48, 49), such that the system remains in local mechanical equilibrium over an extended period of time, as captured by the stable patterns in the Hi-C data (14). Although a great amount of cell-to-cell variation is expected for a population of cells (39, 49, 50), fluorescence measurement indicates that the spatial distances between pairs of chromatin segments can be well described by the gaussian distribution (21, 51–53) (see Fig. S1). This motivates us to model the chromosome structure using a gaussian polymer network whose configuration fluctuates around its local mechanical equilibrium state (54, 55). See SI Appendix for more justifications for the use of the Gaussian polymer network model.

For a polymer chain whose long-range pairwise interactions are restrained via harmonic potentials with varying stiffness, the distance between each pair of segments *i* and *j* is written in the following form

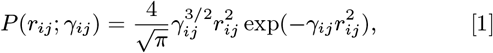

with 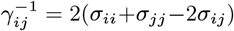, where *σ_ij_* is the positional covariance determined by the topology of polymer network (14, 42). The contact probability between the two segments, *p_ij_*, is the probability that their pairwise distance *r_ij_* is below a cutoff *r_c_*. In other words, it is calculated using 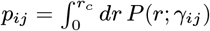, where *P* (*r*; *γ_ij_*) is the distance distribution in Eq. 1. Importantly, this model establishes a one-to-one mapping from the contact probability *p_ij_* to the parameter *γ_ij_*, which allows one to determine the covariance matrix {*σ_ij_*} and consequently the cross-correlation matrix, 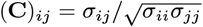 (see Materials and Methods for mathematical details). As a result, the following transformation from *p_ij_* to (**C**)_*ij*_ is conceived:

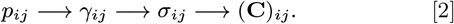

The resulting cross-correlation matrix **C** is used as the input data for our domain inference algorithm (see Fig. 2).

**Fig. 2.**
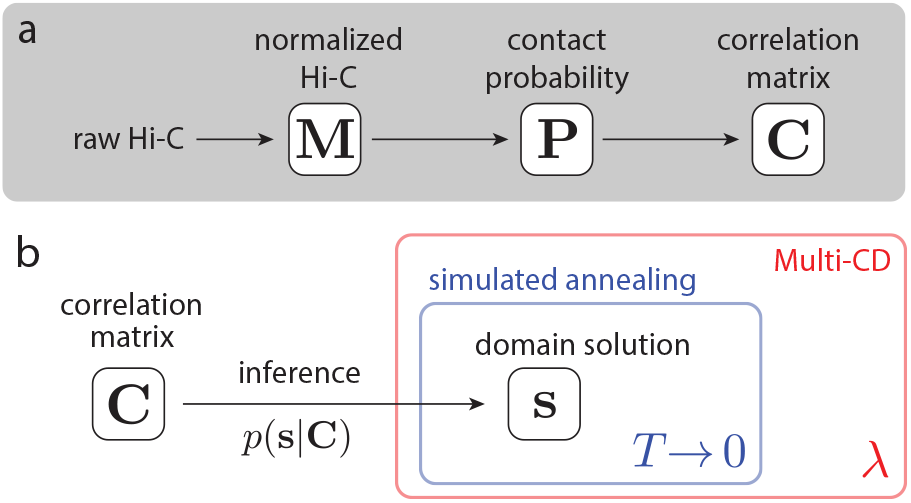
Schematic of the Multi-CD algorithm. **(a)** Pre-processing: Hi-C data provide information of the correlation pattern. **(b)** Inference: for a given correlation matrix **C**, the Multi-CD algorithm finds chromatin domain (CD) solutions **s** at multiple scales. The algorithm of Multi-CD is repeatedly applied to **C** to determine the best domain solutions at different values of *λ* (outer red box). At each *λ*, the best domain solution is found through a simulated annealing, in which the effective temperature *T* is gradually decreased (inner blue box). We use a MCMC sampling method to approximate the posterior distribution 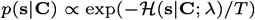 and to find the domain solution at each fixed temperature *T*. The boxes in red and blue represent the two levels of iterations varying *λ* and *T*, respectively, in the algorithm.

We propose a principled interpretation of Hi-C data as a *contact probability matrix*, which can be derived from a mathematically tractable yet physically meaningful model of gaussian polymer network. As a pre-processing method, our approach can replace the common but arbitrary use of nonlinear (most often logarithmic) scaling of the Hi-C data.

### Modeling the correlations with the group model

Clustering a correlation matrix into a finite number of correlated groups is a general problem discussed in diverse disciplines. We adapted a statistical mechanical formalism known as the “group model,” developed for identifying the correlated groups of companies from empirical data of stock market price fluctuations (56–58). Without ambiguity, the formalism can be applied to the clustering of correlated genomic segments in a chromosome.

Let us assume that each genomic segment *i* ∈ {1, 2,···, *N*} belongs to a chromatin domain *s_i_*. Then the vector **s** = (*s*_1_, *s*_2_, …, *s_N_*) can be called the *domain solution* for the *N* segments. For example, a state **s** = (1, 1, 1, 2, 2, 3) describes a structure where the 6 genomic segments are clustered into 3 domains. Indexing of the domains is arbitrary. If there are *K* distinct domains in the solution, we can always index the domains such that *s_i_* ∈ {1, 2,···, *K*}.

We also assume that the cross-correlation matrix **C**, captured by Hi-C, is essentially described by the correlation of a set of hidden variables {*x_i_*} where *x_i_* represents the “genomic state” of the *i*-th chromatin segment. Without loss of generality we can only consider the case where *x_i_* has zero mean and unit variance; or simply standardize as 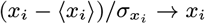. Adapting the formalism in Refs. (56, 57), we assume that each *x_i_* obeys the following stochastic equation

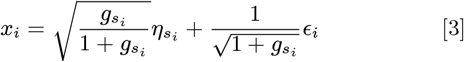

where 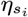 and *ϵ_i_* are two independent and identically distributed (i.i.d) random variables with 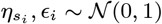, that are linked to the domain (*s_i_*) and the individual segment (*i*) respectively. The parameter 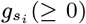 is associated with each domain *s_i_*, such that a larger 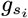 indicates a stronger contribution from the domain-dependent variable 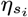. The cross-correlation between two segments *i* and *j* is written as

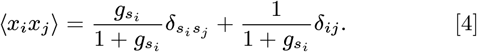

In light of Eq. 4, the first term of Eq. 3 on the right hand side contributes to intra-CD correlation of the *s_i_*-th CD with increasing 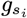; the second term of Eq. 3 corresponds to a noise that randomizes the intra-domain correlation. Regarding (**C**)_*ij*_ calculated from Hi-C as the correlation between the stochastic variables *x_i_* and *x_j_*, each representing the genomic state of segment (Eq. 4), namely,

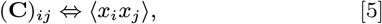

we use **C** as an input data for the clustering approach prescribed by the group model.

Our goal is to find the domain solution **s** = (*s*_1_, *s*_2_, ···, *s_N_*) that best represents the pattern in the correlation matrix **C**, with an appropriate set of strength parameters **g** = (*g*_1_, *g*_2_,···, *g_K_*), where *K* is the number of distinct domains in the solution. Using the group model described above, we can calculate the likelihood *p*(**C**|**s**, **g**) that the observed correlation matrix **C** was drawn from an underlying set of domains **s** with strengths **g** (see SI Appendix for derivation). The log-likelihood is written as a sum over all domains in the solution **s**:

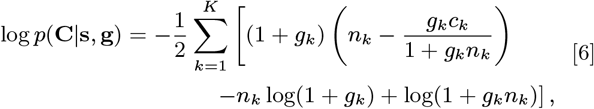

where 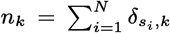 is the size of domain *k*, and 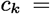 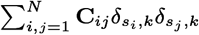 is the sum of all intra-domain correlation elements. The log-likelihood in Eq. 6 is maximized at 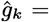 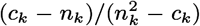 for each *k*, allowing us to consider the reduced likelihood 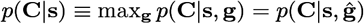.

For convenience, we define an energy-like cost function 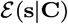 for a domain solution **s** given a correlation matrix **C**, such that the problem of finding the maximum-likelihood solution **s** is equivalent to finding the **s** that minimizes 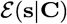. Specifically, it is useful to write the likelihood function as 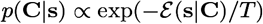 to resemble a Boltzmann distribution, with

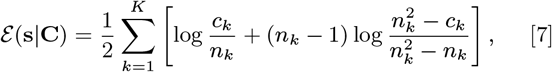

where the new parameter *T* > 0 corresponds to the thermodynamic temperature. The best domain solution can be found by using the simulated annealing method where *T* is slowly decreased.

### Inferring the domain solutions at multiple scales: a tunable group model

Besides evaluating how well a domain solution **s** explains the correlation pattern in the data, which is captured by the likelihood, we also want to impose an additional preference to more parsimonious solutions. This is done by introducing a prior distribution, *p*(**s**). Here we use the form 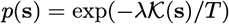, where 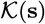 is a function that gives a larger value for a more fragmented solution **s** (larger number of domains), and *λ* (≥ 0) is a parameter that scales the overall strength of the prior. Specifically, 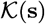 is defined such that 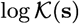 is the entropy of **s**:

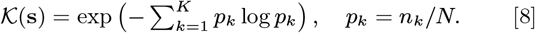

This quantity measures the effective number of domains; for example, 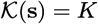 when the domain sizes are uniform.

Taking a Bayesian approach, we combine the likelihood and the prior, to construct the posterior distribution according to the Bayes rule: *p*(**s**|**C**) ∝ *p*(**C**|**s**) *p*(**s**). The posterior is written in terms of an effective Hamiltonian 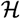:

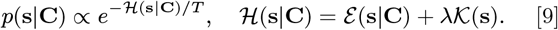

Finding the *maximum a posteriori* (MAP) estimate, **s**\* = argmax_**s**_ *p*(**s**|**C**), is equivalent to solving a global minimization problem for 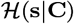. As the **s**-space is expected to be high-dimensional and is likely characterized with multiple local minima, we use simulated annealing (59) to find the energy-minimizing **s**\* (Fig. S2) (see Materials and Methods for the details).

Our algorithm, Multi-CD, is a principled method for inferring the chromatin domain (CD) solutions at multiple scales. The group model becomes “tunable” at the level of inference through the parameter *λ*: as *λ* is increased, the resulting estimate **s**\* tends to have a fewer number of domains. In parallel to the statistical physics problem of a grand-canonical ensemble, *T* is the effective *temperature* of the system, and *λ* amounts to the negative *chemical potential*.

## Results

### Discovery of chromatin domains at multiple scales

Given a raw Hi-C matrix, one can use our Multi-CD algorithm for transforming it to a correlation matrix **C** (Fig. 2a), as well as for identifying a set of CDs for each fixed *λ* (Fig. 2b).

Here we applied Multi-CD on a sample subset of Hi-C data from a commonly used human lymphocyte cell line, GM12878, at 50-kb resolution. After the transformation of the raw Hi-C (Fig. 3a) into a correlation matrix (Fig. 3b), Multi-CD was employed to infer a family of CD solutions that vary with *λ* (Fig. 3c). We also applied Multi-CD to four other cell lines, HUVEC, NHEK, K562, and KBM7 (Fig. 3d), and analyzed the CD solutions for all five cell lines. All results shown in the main text consider chromosome 10; see Fig. S3 for similar results from three other chromosomes (chr4, 11, 19).

**Fig. 3.**
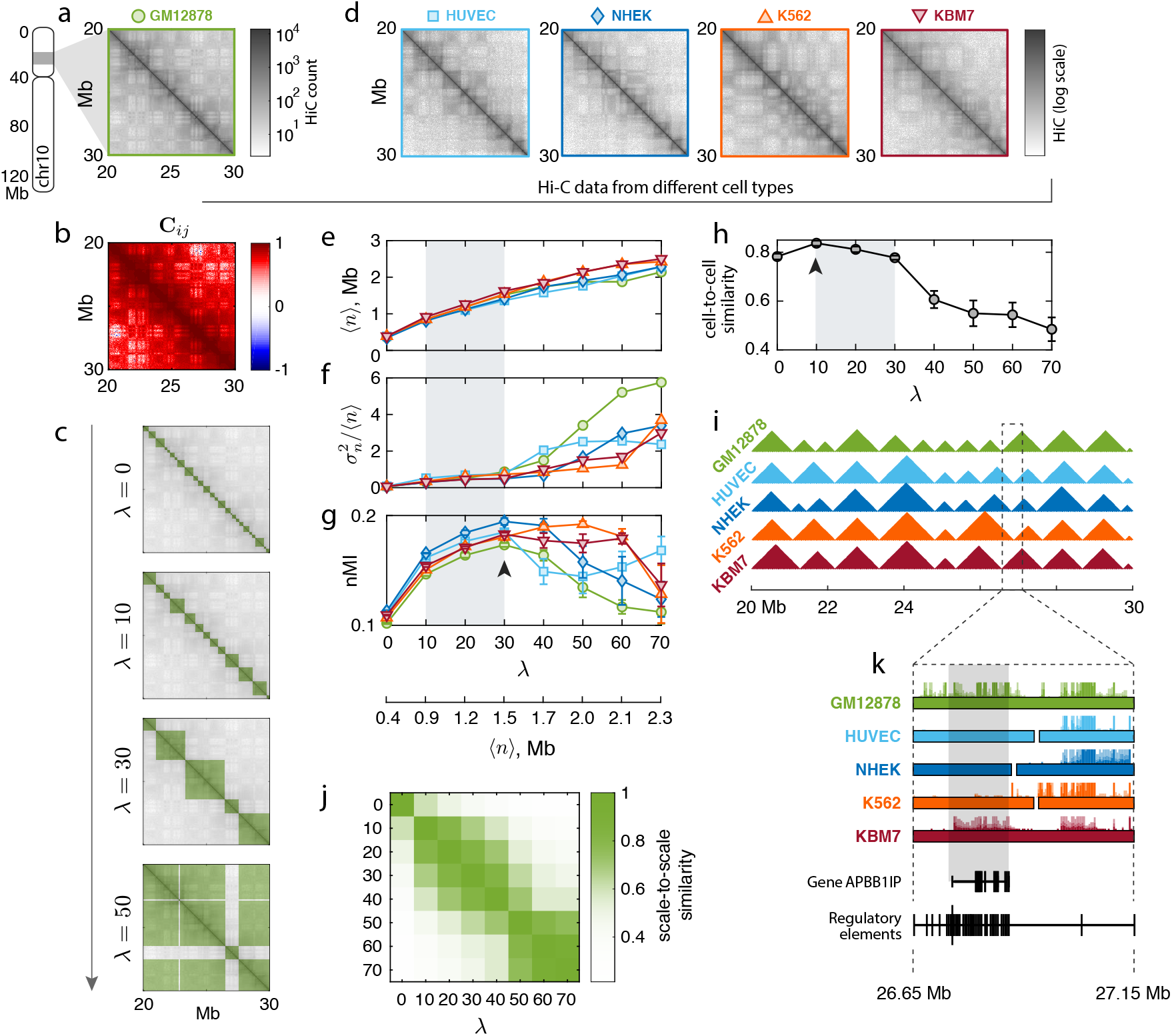
Multi-scale chromatin domain solutions for various cell types. **(a)** A subset of 50-kb resolution Hi-C data, covering a 10-Mb genomic region of chr10 in GM12878. (b) The cross-correlation matrix **C**_*ij*_ for the corresponding subset. See Fig. S4 for a full-chromosome view. **(c)** Multi-CD applied to the correlation matrix in **b**. Domain solutions determined at 4 different values of *λ* = 0, 10, 30, 50. **(d)** Hi-C data from the same chromosome (chr10) in four other cell lines: HUVEC, NHEK, K562, and KBM7. Same subset as in **a**. **(e-g)** Characteristics of the domain solutions determined for all five cell lines in **a** and **d**: **(e)** the average domain size, ⟨*n*⟩; **(f)** the index of dispersion in the domain size, 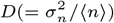; **(g)** the normalized mutual information, nMI. **(h-i)** Comparison of domain solutions across cell types. **(h)** Average cell-to-cell similarity of domain solutions, in terms of Pearson correlations, at varying *λ*. **(i)** Domain solutions obtained at *λ* = 10 for 5 different cell types. See Fig. S5 for solutions at *λ* = 0 and *λ* = 40. **(j)** Similarity between domain solutions at different *λ*’s, shown for GM12878. See Fig. S6 for corresponding results for the other four cell lines. **(k)** RNA-seq signals from the five cell lines (colored hairy lines), on top of the TAD solutions (filled boxes), in a genomic interval that contains the regulatory elements associated with a gene APBB1IP. APBB1IP is transcriptionally active only in two cell lines, GM12878 and KBM7, where the regulatory elements are fully enclosed in the same TAD. See Fig. S7 for a larger figure.

We observed several general features from the families of CD solutions in these cell lines:

i. The average domain size ⟨*n*⟩ always increased monotonically with *λ* (Fig. 3e), as expected from our construction of the prior.
ii. The domain sizes were relatively homogeneous in the small-domain regime (small *λ*), but became heterogeneous after a cross-over point (Fig. 3f). To quantify this, we defined the *index of dispersion* for the domain sizes, which is simply the variance-to-mean ratio 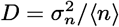. If the domains were generated by randomly selecting the boundaries along the genome, the dispersion would be *D* = 1; a smaller *D* < 1 indicates that domain sizes are more homogeneous than random. A larger *D* > 1 means that the domain sizes are heterogeneous. The crossover points arise at *λ_cr_* ≈ 30 − 40 (⟨*n*⟩_*cr*_ ≈ 1.6 Mb) for GM12878, HUVEC, and NHEK; and *λ*_*cr*_ ≈ 60 − 70 (⟨*n*⟩_*cr*_ ≈ 2.2 Mb) for K562 and KBM7. We observed that the onset of heterogeneity was related to the appearance of non-local domains (Fig. 3c).
iii. We quantified the goodness of each CD solution by comparing its corresponding binary matrix against the Hi-C data in terms of the normalized mutual information (nMI; see Materials and Methods for details). There is a scale, *λ*\*, at which the diagonal block pattern manifested in Hi-C data is most accurately captured. In Fig. 3g, the best solution was found at *λ*\* ≈ 30 for GM12878, HUVEC, and NHEK; the *λ*\*’s were identified at larger values for K562 and KBM7. As an interesting side note, K562 and KBM7 belong to immortalized leukemia cell lines, whereas the other three cell types are normal cells; the different statistical property of Hi-C patterns manifested in *λ*\* may hint at a link between the pathological state and a coarser organization of the chromosome.
iv. The CD solutions inferred by Multi-CD, especially the families of local CDs, appeared to be *conserved* across different cell types (Fig. 3h-i). We quantified the extent of domain conservation, in terms of the Pearson correlation (see Materials and Methods), averaged over all pairs of different cell types. Domain conservation was strong for smaller domains at *λ* ≤ 30 (⟨*n*⟩ ≲ 1.5 Mb), with the strongest conservation at *λ* = 10 (Fig. 3h). The CDs at *λ* = 10 are shown in Fig. 3i for five different cell types.
v. Finally, we quantified the similarity between pairs of CD solutions obtained at different scales, again using the similarity measure based on Pearson correlation. In the case of GM12878, the family of CD solutions is divided into two regimes; the smaller-scale CD solutions from a range of 10 ≤ *λ* ≤ 40 are correlated among themselves, and the larger-scale CD solutions from *λ* > 40 as well. CD solutions below and above *λ* ≈ 40 are not correlated with each other (Fig. 3j; also see Fig. S6 for the other cell lines).

The division boundary in (v) is found at a *λ* value in the similar range with the best-clustering scale *λ*\*, and the crossover *λ_cr_* from local/homogeneous to non-local/heterogeneous CDs (compare Fig. S6 to Fig. 3f-g). Hereafter we will refer to the two regimes as the family of *local* CDs with homogeneous size distribution (*λ* ≲ *λ*\*), and the family of *non-local* CDs with heterogenous size distribution (*λ* ≳ *λ*\*).

### TAD-like organizations in the family of local CDs

We identified at least three important scales in the family of local CDs. First of all, there was a scale at which domain conservation was maximized across different cells (*λ* = 10). This observation is consistent with the widely accepted notion that TADs are the most well-conserved, common organizational and functional unit of chromosomes, across different cell types (27, 60). Thus, for the example from human chromosome 10, we identify the CDs found at this scale *λ* = 10 as the TADs. The average domain size at *λ* = 10 was ⟨*n*⟩ ≈ 0.9 Mb, which agrees with the typical size of TADs as suggested by previous studies (22, 23).

The goodness of clustering, on the other hand, was maximized at a larger scale, *λ* ≈ 30 (⟨*n*⟩ ≈ 1.5 Mb) for GM12878. The CDs at this scale turned out to be aggregates of multiple TADs in the genomic neighborhood, from visual inspection (see Fig. 3c), or as quantified in terms of our nestedness score (see Materials and Methods). We therefore identify these CDs as the “meta-TADs”, a higher-order structure of TADs, adopting the term of Ref. (6). In contrast to a previous analysis that extended the range of meta-TADs to the entire chromosome (6), we use the term meta-TAD exclusively for the larger-scale local CDs, distinguishing them from the non-local structures (i.e., compartments, discussed below). We note, however, that the terminologies of TADs and the meta-TADs are still not definitive – a recently proposed algorithm based on structural entropy minimization (61) found that the “best” solutions were found at ~ 2 Mb domains, which is consistent with our findings, although these domains were called the TADs in Ref. (61).

Finally, a trivial but special scale is *λ* = 0, where no additional preference for coarser CDs is imposed. The CDs at this scale are supposed to best explain the local correlation pattern that is reflected in the strong Hi-C signals near the diagonal. These smaller CDs are almost completely nested in the TADs and the meta-TADs; we can therefore call them the sub-TADs. We also confirmed that the sub-TAD solutions were not limited by the resolution of the Hi-C data; sub-TADs were robustly reproduced from a finer, 5-kb Hi-C (Fig. S8).

The first three panels in Fig. 3c shows three representative TAD-like CD solutions at *λ* ≤ *λ*\*: sub-TADs (*λ* = 0; smallest CDs), TADs (*λ* = 10, strongest domain conservation), and meta-TADs (*λ* = *λ*\* = 30, largest nMI). The nested structure is reminiscent of the hierarchically crumpled structure of chromatin chains (37, 62).

### Chromatin organization and its link to gene expression

The CD solutions from Multi-CD can shed important insights into the link between chromatin organization and gene expression. To demonstrate this, we overlaid the RNA-seq profiles on the TAD solutions, identified for the corresponding subset of the chr10 of five cell lines (GM12878, HUVEC, NHEK, K562, KBM7) (Fig. 3k; also see Fig. S7). At around 26.8 Mb position of this chromosome, we found a gene APBB1IP, which is transcriptionally active in GM12878 and KBM7 but not in HUVEC, NHEK and K562. Consulting the GeneHancer database (63), we identified the regulatory elements for this gene (enhancers and promoters) in the interval between 26.65 and 27.15 Mb. Notably, our Multi-CD solutions show that the interval associated with the regulatory elements is fully enclosed in the same TAD in GM12878 and KBM7, whereas it is split into different TADs in the other three cell lines (Fig. 3k). The observation suggests that, for a gene to be expressed, it is critical that all regulatory elements are within the same TAD; this is consistent with the understanding that TADs define the functional boundaries for genetic interactions (5, 6, 17, 20).

### Compartments as the best domain solution that coexists with TAD-like domains

The super-Mb sized domains are generally defined as the compartments in the chromosome organization (27). Compartments are characterized by the checkerboard pattern in off-diagonal part of correlation matrix, being highly non-local CDs. Our formulation of the group model has the flexibility for dealing with the non-local CDs naturally. However, a naïve application of Multi-CD by increasing *λ* did not identify the compartments; some non-local CDs were found (Fig. 3c), but they do not correspond to alternating patterns characteristic to compartments.

We hypothesize that compartments correspond to a secondary CD solution, that coexists with a best solution that was already identified. Assuming a statistical independence between the two solutions and the additivity of cross-correlations (extension of Eq. 3), the inference of the secondary solution is reduced to a standard Multi-CD applied to a modified input data, which is essentially the result of taking out the best CD solution from the original correlation matrix **C** (see SI Appendix). Here we consider a simplified version of this problem, and remove from **C** a diagonal band of width 2 Mb, similar to the size of meta-TADs (Fig. 4a). Applied to the modified Hi-C, Multi-CD successfully captures the non-local correlations, and identifies two large compartments with alternating patterns (Fig. 4b). The correspondence is clearer when the indices of segments are re-ordered (Fig. 4c). Because the larger CD (*k* = 1) shows a greater number of contacts (Fig. S9), it can be associated with the B-compartment, which is usually more compact; *k* = 2 is associated with the A-compartment. Further validation of the two compartments will be presented below, through comparisons with epigenetic markers.

**Fig. 4.**
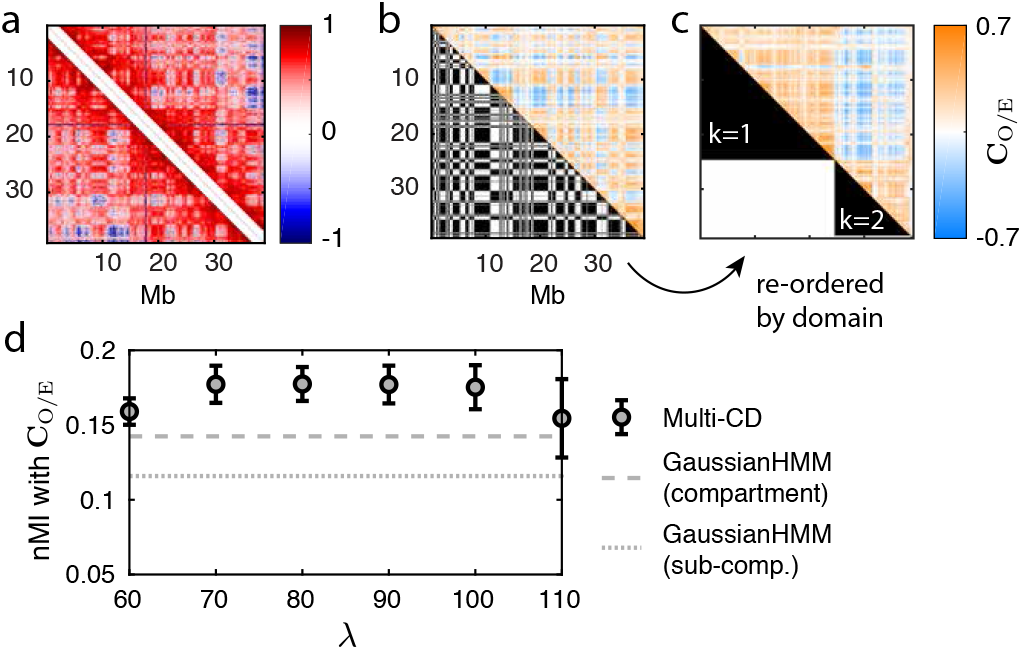
Domain solutions for compartments. **(a)** Input correlation data for compartment identification. The 2-Mb diagonal band was removed. **(b)** Lower triangle: CDs obtained at *λ* = 90, based on the diagonal-band-removed data, which we identify as the compartments. Upper triangle: the **C**_O/E_ matrix shown for comparison. **(c)** Same pair of data, after re-ordering to collect the two largest CDs in our solution (lower triangle), with *k* = 1, 2 as the B- and A-compartments respectively. The **C**_O/E_ is simultaneously reordered to show a clear separation of correlation patterns (upper triangle). **(d)** nMI between CDs at varying *λ* and **C**_O/E_, showing a plateau in the range 70 ≤ *λ* ≤ 100. CDs inferred by Multi-CD show consistently higher nMI, compared to sub-compartments (dashed line) and compartments (dotted line) from a previous method (19).

To compare the goodness of our compartments with existing methods, we calculated the nMI against the **C**_O/E_ matrix, the conventional form for compartment identification (see Materials and Methods) (18). We find that Multi-CD outperforms GaussianHMM (19), a widely accepted benchmark, in capturing the large-scale structures in Hi-C (Fig. 4d).

### Multi-scale, hierarchical organization of chromatin domains

Now that we identified four classes of CD solutions, namely sub-TADs, TADs, meta-TADs and compartments, we examined their hierarchical relationships. Note that these CDs were obtained independently at the respective *λ* values, not through a hierarchical merging. Sub-TADs or TADs are almost always nested inside a meta-TAD, and TADs inside a meta-TAD, whereas there are mismatches between the TAD-like domains and the compartments. We quantified this relationship in terms of a nestedness score *h*, such that *h* = 0 indicates the chance level and *h* = 1 a perfect nestedness (see Materials and Methods), along with a visual comparison of each pair of CD solutions (Fig. 5a). This analysis confirms that the hierarchy between any pair of TAD-like domains (sub-TADs, TADs, and meta-TADs) is appreciably strong. On the other hand, the hierarchical links between the TAD-like domains and compartments are much weaker, which is again consistent with the recent reports that TADs and compartments are organized by different mechanisms (64, 65).

**Fig. 5.**
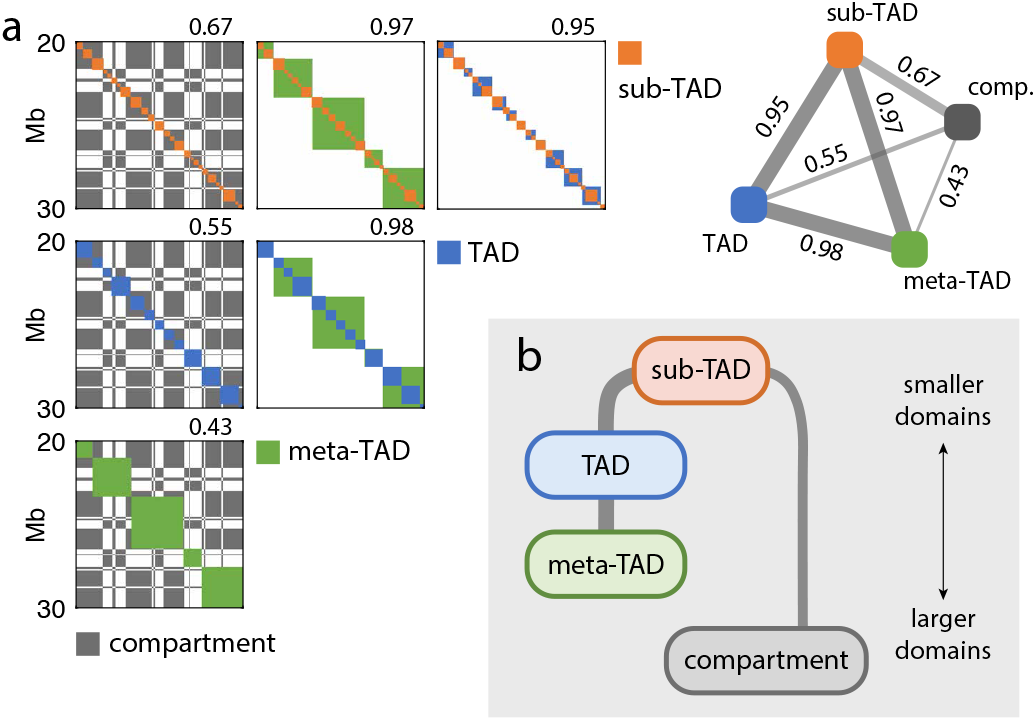
Hierarchical organization of CD families. **(a)** Hierarchical structure of CDs are highlighted with the domain solutions for sub-TADs (red), TADs (green), meta-TADs (blue) and compartments (black). Shown for chr10 of GM12878. Each square panel overlays a pair of CD solutions; number above the panel reports the nestedness score. Inset: a reprint of the nestedness scores in a tetrahedral visualization with the four representative CD solutions. **(b)** A schematic diagram of inferred hierarchical relations between sub-TADs, TADs, meta-TADs and compartments, based on our calculation of nestedness scores.

Although the nestedness score between sub-TADs and compartments (nestedness score *h* = 0.67) is not so large as those among the pairs of TAD-based domains, it is still greater than those between TADs and compartments (*h* = 0.55) or between meta-TADs and compartments (*h* = 0.43). Thus, sub-TAD can be considered a common building block of the chromatin architecture (see Fig. 5b).

### Validation of domain solutions from Multi-CD

The CD solutions from Multi-CD are in good agreement with the results of several existing methods. Specifically, our CDs correspond to the previously proposed sub-TADs (19) at *λ* = 0, to the TADs (22) at *λ* ≈ 10, and to the compartments (19) at *λ* ≈ 90 (see Fig. S10). When assessed in terms of the nMI, Multi-CD outperforms the corresponding alternatives (ArrowHead (19), DomainCaller (22), GaussianHMM (19) for sub-TADs, TADs, meta-TADs) at the respective scales (Fig. 6a).

**Fig. 6.**
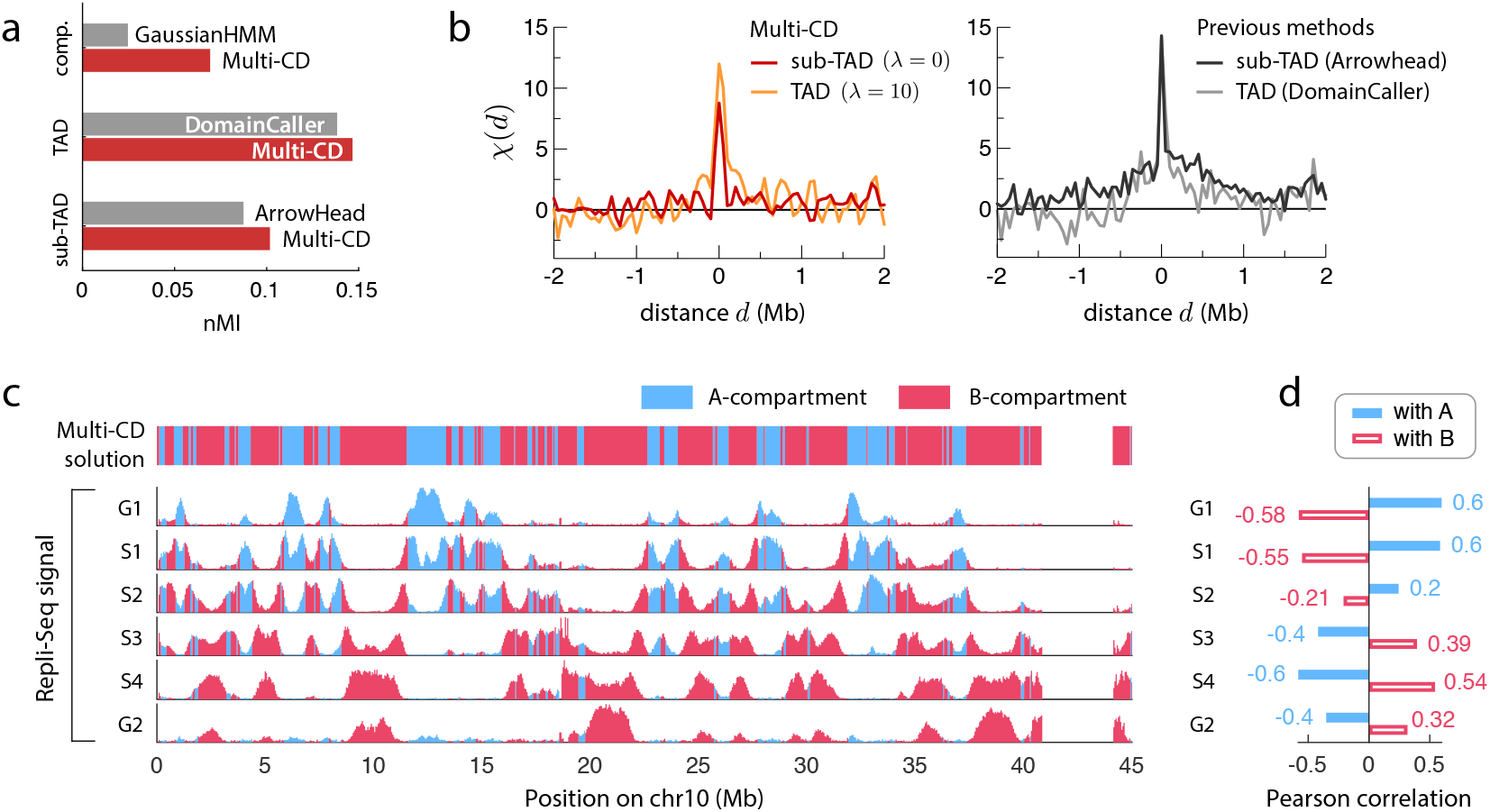
Validation of CD solutions from Multi-CD. **(a)** In terms of the normalized mutual information between the CD solutions and the input data, Multi-CD outperforms ArrowHead, DomainCaller and GaussianHMM at the corresponding scales (sub-TAD, TAD and compartment). **(b)** The correlation function *χ*(*d*) between CTCF signals and the domain boundaries. Shown for sub-TADs and TADs, obtained from Multi-CD (left); from ArrowHead and DomainCaller (right). **(c)** Genome-wide, locus-dependent replication signal. Top panel shows the A- (blue) and B- (red) compartments inferred by Multi-CD. Bottom panels show the replication signals in six different phases in the cell cycle, shaded in matching colors for the two compartments. **(d)** Pearson correlation between the replication signals and the two compartments A (filled blue) and B (open red).

In order to further validate the biological relevance of the CD solutions from Multi-CD, we compared them with several biomarkers that are known to be correlated with the spatial organization of the genome (66). All results shown here are for chr10 of GM12878.

First, we calculated how much the boundaries of our sub-TAD and TAD solutions are correlated with the CTCF signals, which are known to be linked to TAD boundaries (22, 23) (Fig. 6b). We quantified this in terms of a correlation function, *χ*(*d*), where *d* is the genomic distance between a domain boundary and each CTCF signal (see Materials and Methods). The correlation function shows a strong enrichment of CTCF signals at domain boundaries (high peak of correlation at *d* ≈ 0), as well as precision (fast decay of correlation as *d* increases). Multi-CD performs similarly to ArrowHead and DomainCaller in terms of the enrichment at the boundary, and does better in terms of the precision (Fig. 6b). Specifically, when fitted to exponential decays, the correlation lengths are 34 kb (*λ* = 0) and 143 kb (*λ* = 10) for Multi-CD, compared to ≳ 900 kb for the two previous methods (Fig. 6b).

Next, we compared our compartment solutions (CDs at *λ* = 90, shown in Fig. 4b) with the replication timing profiles (Repli-Seq), which are known to correlate differently with the A- and B- compartments (7, 67). Our inferred compartments exhibit the anticipated patterns of replication timing (Fig. 6c); the A-compartment shows an activation of replication signals in the early-phases (G1, S1, S2) and a repression in the later phases (S3, S4, G2), whereas the B-compartment shows an opposite trend. There is a clear anti-correlation between the replication patterns in the two compartments along the replication cycle (Fig. 6c), as quantified in terms of the Pearson correlation (Fig. 6d). Comparison to other epigenetic markers, such as the pattern of histone modifications, further confirms the association of our CD solutions with the A/B-compartments (Fig. S11).

## Discussion

Multi-CD has many essential advantages that will make it a useful tool for the study of chromatin organization. As a computational algorithm, Multi-CD includes two core steps: the pre-processing of raw Hi-C data into a correlation matrix, and the inference of chromatin domain (CD) solutions from the correlation matrix. The pre-processing is based on a model of gaussian polymer network, allowing a physically justifiable interpretation of the Hi-C data.

The domain identification problem is formulated by combining the group model with a Bayesian inference for the CD solution. The formulation of Multi-CD that optimizes the recognition of global pattern appeared in Hi-C naturally deals with non-local CDs, which differentiates Multi-CD from previous methods that focus on local features in Hi-C, such as CD boundaries or loops enriched with higher contact frequencies. Moreover, Multi-CD can find CDs across a wide range of scales without having to adjust or down-sample the Hi-C data to match the scale of CDs to be identified, which is an important improvement over many existing methods.

An important feature of Multi-CD, as emphasized in the name, is that it provides a unified framework to identify CDs at multiple scales, where the scales of the CDs are tuned by a single parameter *λ*. The resulting *family* of CD solutions allow quantitative comparisons between CD solutions at different scales. The analysis revealed special scales at which the CD solutions are particularly interesting: sub-TADs (*λ* = 0), TADs (*λ* = 10, where domain conservation was strongest), and meta-TADs (*λ* = 30, where the correlation pattern was best captured). At larger scales, we found that compartments (*λ* = 90) emerge as a secondary solution that can be inferred after removing the local signals in the correlation matrix that correspond to the TAD-like solutions. We confirmed that Multi-CD successfully reproduces, or even outperforms, the existing methods to identify CDs at the specific scales. Importantly, Multi-CD achieves this performance through a single unified algorithm, which not only identifies the specific CD solutions accurately, but also allows a comparative analysis of the multi-scale family of solutions.

In particular, we characterized the hierarchical organization of the chromatin by quantifying the similarity and the nestedness between CD solutions at two different scales. We showed that the characteristics of CD solutions shared by the local, TAD-like domains do not precisely hold together in the non-local, compartment-like domains. This finding is consistent with the recent studies which report that compartments and TADs are formed by different mechanisms of motor-driven active loop extrusion and microphase separation, and that they do not necessarily have a hierarchical relationship (65, 68–70). Meanwhile, the sub-TADs are nested in each of the other three solutions, including the compartments (Fig. 5), indicating that sub-TADs are the fundamental building blocks of the higherorder CD organization. In fact, the existence of sub-TADs is robust when a finer-resolution Hi-C is considered. Applying Multi-CD on Hi-C data at 5-kb resolution, we clearly recover the sub-TADs that are consistent with the sub-TADs obtained from the 50-kb Hi-C (see Fig. S8).

While there are methods that report hierarchical CDs (32, 33), Multi-CD makes significant advances both algorithmically and conceptually. Multi-CD can detect non-local domains with better flexibility instead of finding a set of intervals. Multi-CD also avoids the high false-negative rate that is typical of the previous method (e.g., TADtree (32)) that focuses on the nested domain structure (Fig. S12). Further, employing an appropriate prior to explore the solution space effectively, Multi-CD can avoid the problem encountered in Armatus (33) which skips detection of domains in some part of Hi-C data while its single scale parameter is varied (Fig. S12).

Multi-CD is a method of great flexibility that can be readily applied to analyze any dataset that exhibits pairwise correlation patterns. However, two cautionary remarks are in place for more careful interpretation of the results. (i) In general, the relevant values of *λ* depend on the resolution of the input Hi-C dataset, as well as on the cell type. While *λ* is a useful parameter that allows comparative analysis, its specific value does not carry any biological significance. Although we referred to a specific CD solution by the corresponding value of *λ* in the current analysis (Fig. 3), the lesson should *not* be that TADs, for instance, always correspond to the particular value of *λ*; instead, TADs should be identified as the most conserved CD solutions across cell types after scanning a range of *λ*’s. (ii) Multi-CD is agnostic about whether the collected data is homogeneous or heterogeneous. Application of Multi-CD to single-cell Hi-C data, and the subsequent interpretation of the result, would be straightforward; however, if the input Hi-C data were an outcome of a mixture of heterogeneous subpopulations, the solution from Multi-CD would correspond to their superposition. This is a fundamental issue inherent to any Hi-C data analysis method. Despite the presence of cell-to-cell variations, the population-averaged pattern manifest in Hi-C carries a rich set of information that is specific to the cell type. The need for interpretable inference methods that can extract valuable insights into the spatial organization of the genome, including ours, is still high.

To recapitulate, in order to glean genome function from Hi-C data that varies with the genomic state (10–13), a computationally accurate method to identify CD structures is of vital importance. Multi-CD is a physically principled method that identifies multi-scale structures of chromatin domains by solving the global optimization problem. We find the chromatin domains identified from Multi-CD in excellent match with biological data such as CTCF binding sites and replication timing signal. Quantitative analyses of CD structures identified across multiple genomic scales and various cell types offer general physical insight into chromatin organization inside cell nuclei.

## Materials and Methods

### Interpretation of Hi-C data

#### Normalization and contact probability

Here we describe how the Hi-C data can be interpreted as a set of contact probabilities for pairs of genomic segments, *p_ij_*. Typically, a Hi-C matrix have widely varying row-sums; for example, the net count of the *i*-th segment in the experiment is much larger than the net count of the *j*-th segment. To marginalize out this site-wise variation and only focus on the differential strengths of pairwise interactions, the raw Hi-C matrix **M**_raw_ is normalized to have uniform row and column sums. This is achieved using the Knight-Ruiz (KR) algorithm (71), which finds a vector **v** = (*v*_1_,···, *v_N_*) for calculating (**M**)_*ij*_ = *v*_*i*_*v*_*j*_ (**M**_raw_)_*ij*_, such that each row (column) in **M** sums to 1.

We assume that the normalized Hi-C signal is proportional to the contact probability: (**M**)_*ij*_ ∝ *p_ij_*. Note that *p_ij_* is the probability that the two segments *i* and *j* are within a contact distance, and the rows of the contact probability matrix (**P**)_*ij*_ = *p_ij_* is not required to sum to 1. Because the proportionality constant is unknown *a priori*, however, we have a free parameter to choose. We do this by fixing the average nearest-neighbor contact probability, 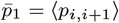. We expect the 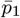 to be relatively close to 1, assuming that nearest-neighbor contact is likely; but not exactly 1, because there are variations among the nearest-neighbor Hi-C signal. In this work we chose 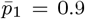. The resulting contact probability matrix **P** is given as 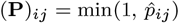, with 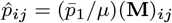, where *μ* = ⟨**M**_*i,i*+1_⟩ is the Hi-C signals averaged over the nearest-neighbors. At 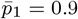, in our case, the fraction of over-saturated elements 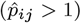 was sufficiently small.

#### Building correlation matrix from Hi-C

A chromosome can be regarded as a polymer chain containing *N* monomers, each of which corresponds to the *i*-th genomic segment and its spatial position is written as **r**_*i*_. Adapting the random loop model (RLM) (42), we interpret chromosome conformation as described by an ideal polymer network, with cross-links between spatially close segments. In RLM, we describe the spatial positions of the polymer segments using a gaussian distribution with zero mean and and a covariance matrix Σ, with elements (Σ)_*ij*_ = *σ_ij_* = ⟨*δ***r**_*i*_ · *δ***r**_*j*_⟩. It follows that the distance *r_ij_* = |**r**_*i*_ − **r**_*j*_| between two monomers *i* and *j* can be described in the form of a weighted gaussian function (Eq. 1) where the variance (2*γ_ij_*)^−1^ is associated with the covariance matrix elements as 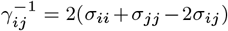. The contact probabilities can be calculated from the distribution of pairwise distances (Eq. 1), by saying that two segments *i* and *j* are in contact when their distance *r_ij_* is below a cutoff, *r_c_*. In other words, we write

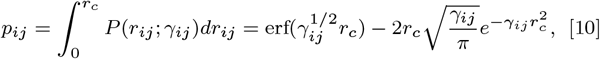

where 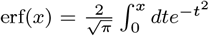. The value of *γ*_*ij*_ is uniquely determined for each *p_ij_*; once we have the *γ_ij_*’s, we can reconstruct the covariance matrix {*σ_ij_*}. Because the diagonals are underdetermined in Hi-C (self-contacts are not reported), we assume a uniform variance *σ_ii_* = *σ_jj_* = *σ_c_* along the diagonal. Note that although the value of *γ_ij_* depends on the choice of *r_c_*, its effect is only to scale the *γ_ij_*’s as 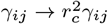, and consequently the *σ_ij_*’s.

Finally, we normalize the covariance matrix to build the correlation matrix **C**:

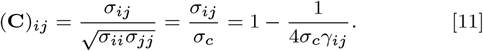

It appears that *σ_c_* sets the overall intensity of **C**. Here, we chose the value of *σ_c_* as the median of 1/4*γ_ij_*, i.e., *σ_c_* = median(1/4*γ_ij_*). This cancels out the scaling effect of *r_c_* in *σ_ij_*, so that the choice of *rc* does not affect the ultimate construction of the correlation matrix **C**.

#### The observed/expected (O/E) matrix and its Pearson correlation matrix

The O/E matrix was used to account for the genomic distance-dependent contact number due to random polymer interactions in chromosome (18). Each pair (*i, j*) in O/E matrix is calculated by taking the count number *M_ij_* (observed number) and dividing it by average contacts within the same genomic distance *d* = |*i − j*| (expected number). Since the expected number could be noisy, one smooths it out by increasing the window size (see refs (18, 19) for further details). In this study, we used the expected number obtained from (19). The Pearson correlation matrix of the O/E (**C**_O/E_) represents the overall contact pattern through pairwise correlation coefficients between segments.

### Determination of optimal CD solutions

#### Metropolis-Hastings sampling

Markov chain Monte Carlo (MCMC) sampling was employed to find the minimum value of the total cost function 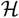. At each trial move from the current state **s** to the next state **s**′, the move is accepted with a probability min(1, *α*), where 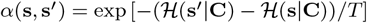.

In sampling the space of CD solutions, a move from a state **s** to another state **s**′ is defined such that the two CD solutions (**s**, **s**′) differ only by one genomic segment. More precisely, because a CD solution is invariant upon permutations of the domain indices, the distance between **s** and **s**′ is uniquely defined as the *minimal* number of mismatches over all possible domain index permutations.

To ensure that the sampling is properly conducted, we continue the sampling until each chain collects *t*_tot_ ≥ 5*τ*\* samples in the CD solution space. Here *τ*\* is the “relaxation time” defined as the number of steps it takes until the autocorrelation function *R*(*τ*), drops significantly: *τ*\* = argmin_*τ*_ |*R*(*τ*) − 1*/e*|. The autocorrelation function is calculated from the value of the total cost function 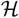, as

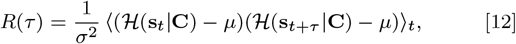

where **s**_*t*_ is the *t*-th sample in the chain, and *μ* and *σ* are the mean and standard deviation of 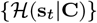, respectively. In Eq.12, the running time average is taken over all the pairs of samples with the time gap of *τ*.

We note that sampling is the computational bottleneck for our method; our stop condition (at 5*τ*\*) was chosen conservatively for accurate solutions. In practice, the sampling time can be reduced at the cost of an increased batch size (number of different initial configurations, as described below in simulated annealing), which is significantly cheaper if parallel computing is used.

#### Simulated annealing

The simulated annealing process is described below.

##### Initialization

An initial configuration **s**^(0)^ is generated in two random steps. First, the total number of CDs, *K*, is drawn randomly from the set of integers {1,···, *N*}. Then, each genomic segment *i* ∈ {1,···, *N*} is allocated randomly into one of the CDs, *k* ∈ {1, 2,···, *K*}. The initial temperature *T*_0_ is determined such that the acceptance probability for the “worst” move around **s**^(0)^ is 0.5.

##### Iteration

At each step *r*, the temperature is fixed at *T_r_*. We sample the target distribution 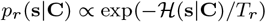, using the Metropolis-Hastings sampler described above. For the next step *r* + 1, the temperature is lowered by a constant cooling factor *c*_cool_ ∈ (0, 1), such that the next temperature is *T*_*r*+1_ = *c*_cool_ · *T_r_*. We used *c*_cool_ = 0.95 in this study.

##### Final solution

The annealing is repeated until the temperature reaches *T_f_*. We used *T_f_* = 0.03. Then we quench the system to the closest local minimum by performing gradient descent. Because there is still no guarantee that the global minimum is found, we tried a batch of at least 10 different initial configurations and chose the final state **s**\* that gives the minimal 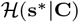.

#### Analysis on subsets of Hi-C data

Our method allows the user to break down the Hi-C data into subsets, as long as the CDs are localized within the subsets (Fig. S13). This saves the algorithm from the large memory requirement of dealing with the entire intrachromosomal Hi-C (for example, Hi-C of chromosome 10 has 2711 bins in 50-kb resolution). For the analysis of the 50-kb resolution Hi-C data in this paper, we used subsets of the data that correspond to 40-Mb ranges along the genome, or 800 bins.

### Analysis and evaluation of domain solutions

#### Similarity of two distinct CD solutions using Pearson correlation

To measure the extent of similarity between two CD solutions **s** and **s**′, we evaluate the Pearson correlation. The binary matrices **B** and **B**′ that represent the two CD solutions, are defined such that the matrix element are all 1’s within the same CD and 0 otherwise. i.e., 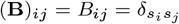. The similarity between **B** and **B**′ is quantified using the Pearson correlation

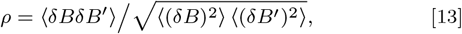

where 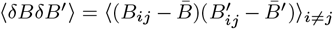, and 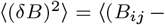 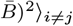. The average ⟨·⟩_*i*≠*j*_ runs over all distinct pairs.

#### Normalized mutual information

We use the mutual information to evaluate how well a CD solution **s** captures the visible patterns in the pairwise correlation data. We consider the binary grouping matrix 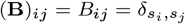 for the CD solution of interest, and compare it to the input data matrix (**A**)_*ij*_ = *A_ij_*. In this study, either log_10_ **M** or **C**_O/E_ was used for **A**. Treating the matrix elements *a* ∈ **A** and *b* ∈ **B** as two random variables, we construct the joint distribution

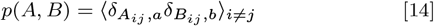

where ⟨·⟩_*i*≠*j*_ is an average over all distinct pairs. The Kronecker delta for the continuous variable *a* is defined in a discretized fashion: that is, 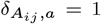 if *A_ij_* ∈ [*a, a* + Δ*a*) and 0 otherwise, where ∆*a*(= [max{*A_ij_*} − min{*A_ij_*}] /100) is used for discretization into 100 bins. Then we can calculate the mutual information,

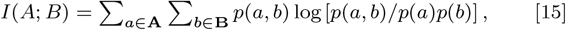

and the noLrmalizeLd mutual information (nMI),

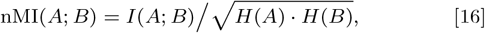

where *H*(*X*) = −∑_*x*∈**X**_ *p*(*x*) log *p*(*x*) is the marginal entropy.

#### Nestedness of CD solutions

Here we define a measure to quantify the nestedness between two CD solutions, **s** (assumed to have smaller domains on average) and **s**′ (larger domains). The idea is the following: **s** is perfectly nested in **s**′ if, whenever two sites belong to a same domain in **s**, they also belong to a same domain in **s**′. For each domain *k* ∈ **s**, we consider the best overlap of this domain *k* on the other solution **s**′:

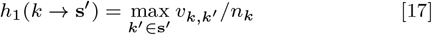

where *v_k,k′_* is the number of overlapping sites between two domains *k* ∈ **s** and *k*′ ∈ **s′**, and *n_k_* is the size of domain *k*. The highest score *h*_1_(*k* → **s**′) = 1 is obtained when domain *k* is fully included in one of the domains in **s**′. The null hypothesis corresponds to where the domains in **s** and **s**′ are completely uncorrelated, in which case *h*_1_ only reflects the overlap “by chance”. The chance level 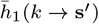 is calculated by making *n_k_* random draws from **s**′; we averaged over 100 independent trials. We normalize the score as

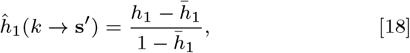

such that 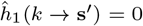 indicates the chance level, and 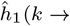 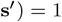 means a perfect nestedness. Finally, we define the nestedness score *h*(**s** → **s′**) for the entire CD solution as a weighted average:

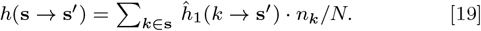

#### Correlation between CTCF signal and domain boundaries

The validity of domain boundaries, determined from various CD-identification methods including Multi-CD, is assessed in terms of their correlation with the CTCF signal. Suppose that the CTCF signal at genomic segment *i* is given as *ϕ*_CTCF_(*i*). Then, we can consider an overlap function between *ϕ*_CTCF_(*i*) and a CD-boundary indicating function *ψ*_DB_(*i*), where *ψ*_DB_(*i*) = 1 if the *i*-th segment is precisely at the domain boundary; *ψ*_DB_(*i*) = 0, otherwise. We evaluated a distance-dependent, normalized overlap function *χ*(*d*), defined as

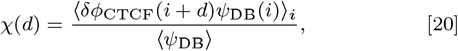

where *δϕ*_CTCF_ = *ϕ*_CTCF_ − ⟨*ϕ*_CTCF_⟩. If the domain boundaries determined from Multi-CD is well correlated with TAD-capturing CTCF signal, a sharply peaked and large amplitude overlap function (*χ*(*d*)) is expected at *d* = 0.

#### Correlation between epigenetic marks and compartments

We calculate the correlation of our compartment solutions with the epigenetic marks. Given a compartment solution **s** with two large domains A and B, we consider two binary vectors **q**(*A*) and **q**(*B*), where 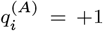 if the *i*-th segment belongs to compartment A, and *q*^(*A*)^ = −1 otherwise. For a set of epigenetic marks measured across the genome is represented with **h**, where its component *h_i_* denotes the value at the *i*-th genomic segment, the correlation between the solutions of compartment *A* and *B* and the epigenetic marks can be evaluated using the Pearson correlations as:

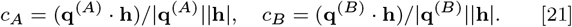

## Data availability

All data used in the paper were obtained from publicly available repositories. See SI Appendix.

## Code availability

The Matlab software package and associated documentation are available online (https://github.com/multi-cd).

## ACKNOWLEDGMENTS

We thank Roger Oria Fernandez for critical comment on our script deposited in GitHub. This work was supported in part by a KIAS Individual Grant at Korea Institute for Advanced Study (No. CG035003 to C.H.). We thank the Center for Advanced Computation in KIAS for providing computing resources.

## Supporting Information Appendix

### A. Data acquisition

#### Hi-C data

Hi-C data were obtained through GEO data repository (GSE63525-*celltype*-primary) (19), where *celltype* is replaced by one of the five different cell types that were considered in our analysis (GM12878, HUVEC, NHEK, K562, and KBM7).

#### Biological markers

The domain solutions from Multi-CD were compared with known biological markers. We obtained these data mostly from the ENCODE project (72). Specifically, we used the enrichment data of the transcriptional repressor CTCF measured in a Chip-Seq assay from http://hgdownload.cse.ucsc.edu/goldenPath/hg19/encodeDCC/wgEncodeUwTfbs/wgEncodeUwTfbsGm12878CtcfStdPkRep1.narrowPeak.gz. We binned the CTCF assay at 50-kb resolution (to match the Hi-C format). If there are multiple signal enrichments in a single bin, we took the average value. Because each CTCF signal has a finite width, there are occasional cases where a signal ranges across two bins; in those cases we evenly divided the signal strength into the two bins. The Repli-seq signals in the six phases G1, S1, S2, S3, S4, and G2 were obtained from http://hgdownload.cse.ucsc.edu/goldenPath/hg19/encodeDCC/wgEncodeUwRepliSeq/, and were averaged over 50-kb windows along the genome to construct the replication timing profiles. The 11 histone mark signals were obtained from http://ftp.ebi.ac.uk/pub/databases/ensembl/encode/integration_data_jan2011/byDataType/signal/jan2011/bigwig/, and undergone the same preprocess as Repli-seq. The RNA-seq data for the four cell lines GM12878, HUVEC, NHEK and K562 were also obtained from http://hgdownload.cse.ucsc.edu/goldenpath/hg19/encodeDCC/wgEncodeCaltechRnaSeq/. RNA-seq for the cell line KBM7 were separately obtained from https://opendata.cemm.at/barlowlab/2015_Kornienko_et_al/hg19/AK_KBM7_2_WT_SN.F.bw. Information about the APBB1IP gene was obtained from the GeneCards database https://www.genecards.org/cgi-bin/carddisp.pl?gene=APBB1IP. The information about its regulatory elements was obtained specifically from the GeneHancer (63) database http://hgdownload.cse.ucsc.edu/gbdb/hg19/geneHancer/geneHancerInteractionsAll.hg19.bb, accessible through the UCSC Genome Browser Module.

### B. Justifications for the Gaussian polymer network for modeling chromosomes

Here we provide additional justifications for the use of harmonic potentials in the effective Hamiltonian, and consequently, a gaussian distribution for pairwise distances (Eq. 1).

The concept of an effective Hamiltonian consisting of harmonic potential terms is not new; it has been widely employed to study a variety of systems, including the phase transition of vulcanized macromolecules with increasing numbers of crosslinks (73, 74), and the fluctuation dynamics of native proteins (gaussian network model, (75)). Furthermore, a slightly modified, but essentially identical, form of Hamiltonian was used to study the dynamics of folding/unfolding transitions of a single RNA molecule under external force (generalized Rouse model, (76)).

Whereas the success of the gaussian polymer network model does not necessarily guarantee its extension to the modeling of chromosomes, our use of a gaussian distribution for the pairwise distance between two segments in the polymer is empirically justified. The Gaussian-like pairwise distance distributions reported by fluorescence measurements of the chromosome (Fig. S1), and the agreement of the 3D structural properties inferred by a modeling approach that shares the same philosophy (*heterogeneous loop model* or HLM, (14)), suggest that a harmonic spring network provides a reasonable approximation of the energy landscape for the mixture of those subpopulations.

As a side note, it is worth highlighting the versatility of the Gaussian polymer network model in representing the complex topology of chromosome conformation. For the conventional Rouse chain whose monomers along the backbone are constrained by an energy hamiltonian 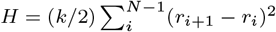 with a uniform spring constant *k*, it is straightforward to show that 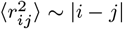. Furthermore, if two monomers are in close proximity to form a contact (*r_ij_* < *r_c_*), then one can obtain the contact probability between monomers *i* and *j* in the chain backbone as 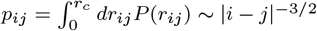. However, adding just a few non-nearest-neighbor interactions to the Rouse model makes the results highly nontrivial. To illustrate this, we explicitly compared the contact probability map of a linear Gaussian chain (Rouse chain), and those of Gaussian polymer network models with varying numbers of non-nearest-neighbor interactions, which were calculated from the HLM-generated structural ensemble (14) (see Fig. S14). The statistical behavior of Gaussian polymer network model differs from that of the linear “Gaussian” chain. The mean square distance 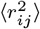 no longer scales linearly with the separation *s* ≡ |*i* − *j*| (see Fig. S14e), and the contact probability *p_ij_* (or *p*(*s*)) is no longer described with a simple scaling relation (Fig. S14f). The simple modification to the Rouse model, resulting in the Gaussian polymer network model, allows one to explore many different issues of chromosomes. In fact, our recent work based on HLM (14) demonstrated several case studies, substantiating the various experimental measurements on chromosome conformation by solving the inverse-problem of inferring chromosome structures from Hi-C data.

### C. Derivation of the likelihood function

Here we show how to derive the likelihood function, Eq. 6.

#### Problem

We want to compute

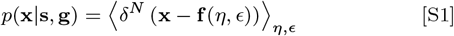

with the following assumptions:

- **x** ∈ ℝ^*N*^ is a sequence of normalized and uncorrelated observations, with zero mean ⟨**x**⟩ = **0**_*N*_ and unit covariance Cov(**x**) = *I_N_*.
- **s** = (*s*_1_, ···, *s_N_*) is a clustering map that assigns each site *i* ∈ {1,···, *N*} to a cluster index *s_i_* ∈ {1,···, *K*}. Without loss of generality, we can assume that *s_i_* ≤ *s_j_* whenever *i* < *j* (ordered indexing).
- 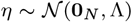 and 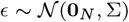 are i.i.d. gaussian random variables, where Λ and Σ are *N* × *N* covariance matrices. The cluster-dependent covariance is a block diagonal matrix 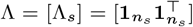, defined element-wise as 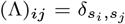. The site-wise variation is assumed to be uncorrelated, with a unit covariance matrix Σ = *I_N_*, or (Σ)_*ij*_ = *δ_ij_*.
- The clustering strength **g** = (*g*_1_,···, *g_K_*) parameterizes the target function **f**, defined element-wise as

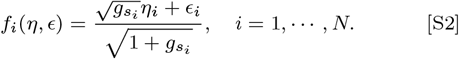

#### Lemma: gaussian integral

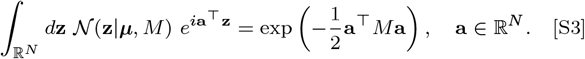

#### Lemma: Sherman-Morrison formula

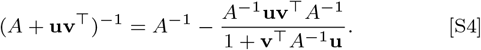

#### Solution

Let us abbreviate the coefficients as 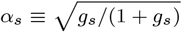 and 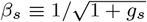, such that 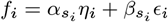. Further define 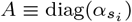 and 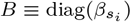, to write **f** = *A**η*** + *B**ϵ***. Taking the inverse Fourier transform of the Dirac delta function, we can write

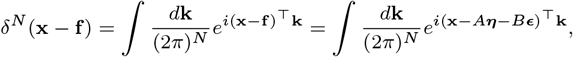

where 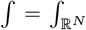 unless otherwise specified. Now we can rewrite Eq. S1, and evaluate the gaussian integrals using the lemma (Eq. S3):

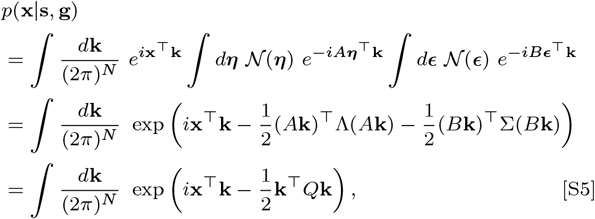

where *Q* ≡ (*A*Λ*A* + *B*Σ*B*). Recognizing that this is another (unnormalized) gaussian integral with covariance matrix *Q*^−1^, use the lemma (Eq. S3) once again:

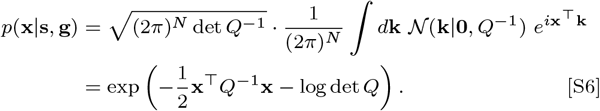

With uncorrelated *ϵ* both *Q* and *Q*^−1^ are block diagonal matrices, the exponent is completely separable by clusters:

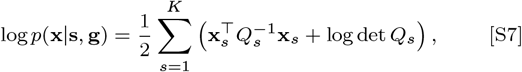

where **x**_*s*_ is the corresponding *n_s_*-dimensional subset of **x**, and 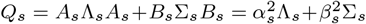, is the *n*_*s*_ × *n*_*s*_ block matrix corresponding to cluster index *s*; element-wise, 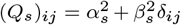.

We now simplify the two terms in the summand of Eq. S7, and show that the resulting expression corresponds to Eq. 6. First, the quadratic term can be expanded by using the Sherman-Morrison lemma Eq. S4:

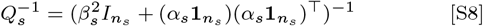

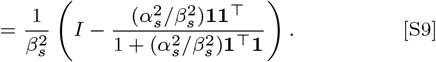

The quadratic form is

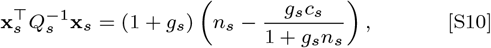

where 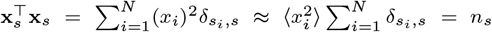, and 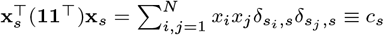.

Second, the log-determinant term can be calculated by considering the eigenvalues of the matrix *Q_s_*. Solving for *Q_s_***z** = *λ_s_***z** for an arbitrary *n_s_*-dimensional vector **z**,

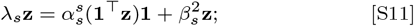

there are two types of solutions. The first possibility is to have the eigenvector **z** ∝ **1**, in which case 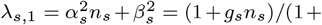 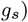. The other possibility is to have 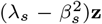 vanish, where 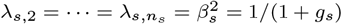; the degenerate eigenvectors span the remaining (*n_s_* − 1)-dimensional subspace. Therefore

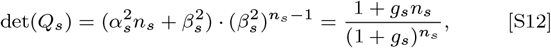

and

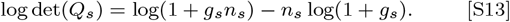

Substituting Eq. S10 and Eq. S13 into Eq. S7, we have derived Eq. 6 as shown the main text.

### D. Modification of the correlation matrix for the secondary domain solution

Here we show how the correlation matrix can be modified to solve for a secondary domain solution (for example compartments), by taking out the contribution from the primary domain solution (for example the meta-TADs), which is supposed to be known already.

We extend the group model to consider a bivariate grouping, *i* ⟼ (*s_i_*, *u_i_*), where each genomic locus *i* simultaneously belongs to a primary group *s_i_* and a secondary group *u_i_*, presumably at different scales. Generalizing Eq. 3, we assume a linear model

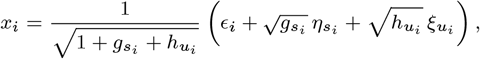

where 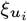 and 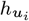 are respectively the random variable and the grouping strength parameter that correspond to the secondary group *u_i_*. If we further assume that *s_i_* and *u_i_* are statistically independent, the pairwise correlation between two loci *i* and *j* can be written as

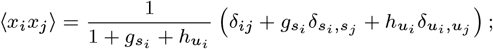

the contributions from different groups are additive.

We can do a straightforward algebra to rearrange the above model as

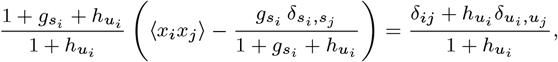

such that the right-hand side becomes the single-group model. The left-hand side of this expression is a normalized residual of the correlation, which we will call 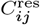. If we have already inferred the primary group *s_i_* and the corresponding strength 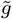 without considering the secondary group, the correspondence to this two-group model (due to normalization) is given as 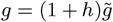. Substituting this, and replacing the model correlation ⟨*x*_*i*_*x*_*j*_⟩ with the data *C_ij_*, the left-hand side of the previous expression is simplified to

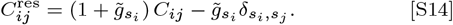

*C*^res^ is written in terms of the original data *C* and the primary solution *s* (and 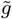) only; it is independent of the unknown secondary solution *u* (and *h*). Now *u* is the solution of a modified single-group problem, using this residual correlation *C*^res^ as the input data.

---

**Fig. S1.**
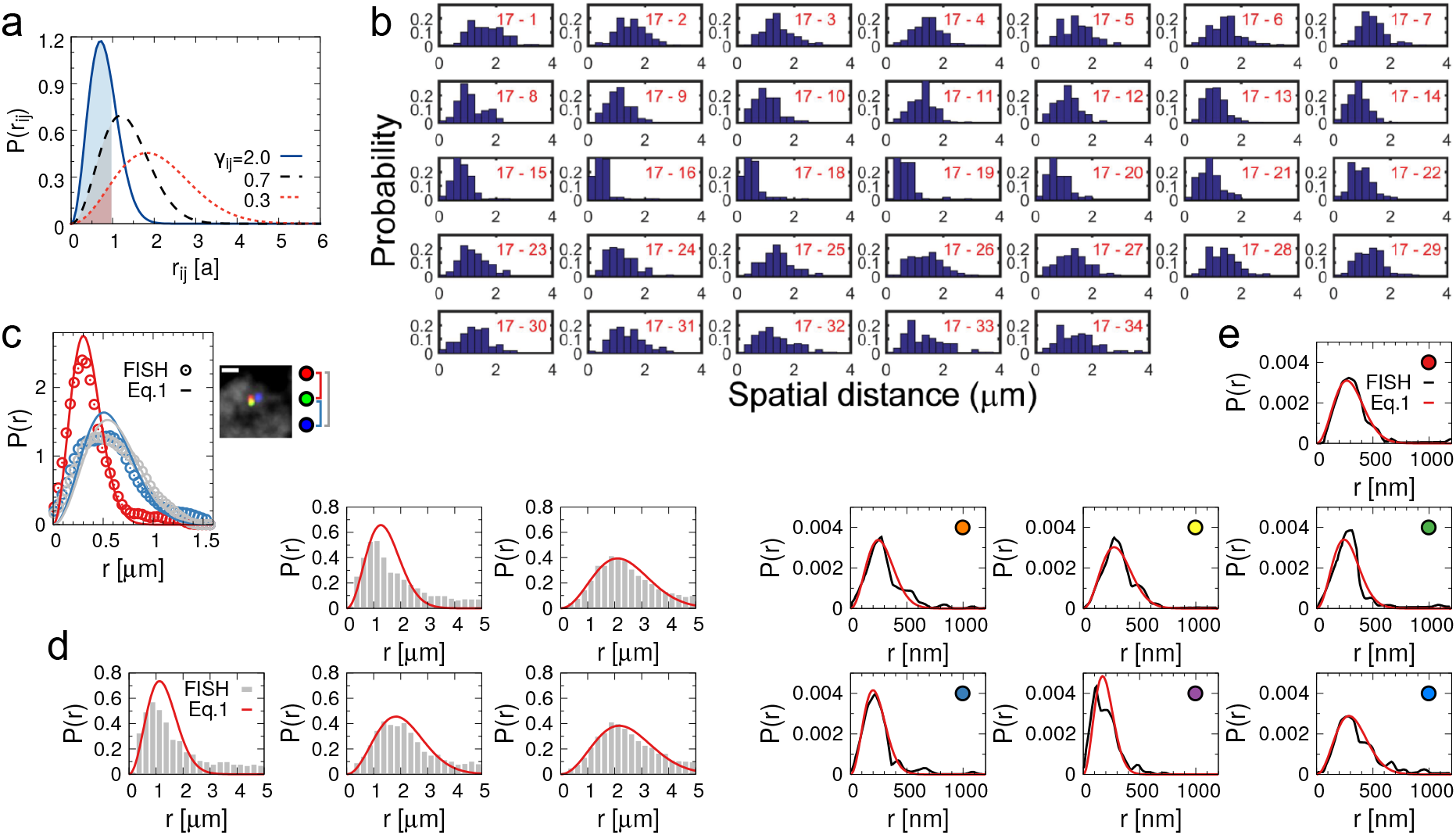
Distance distributions of segment pairs are described by Gaussian. **(a)** Gaussian probability distribution plotting *P* (*r_ij_*) with different values of *γ_ij_* (Eq. 1). The shaded area in different colors represents the corresponding values of contact probabilities, (Eq. 10 at *r_c_* = 1) **(b)** Distance distributions between one TAD (TAD17) and other TADs on Chr21 in human IMR90 cells measured with FISH. This figure was adapted from Fig.S3 in (21). **(c)** Distance distributions between three FISH probes on the X chromosome of male *Drosophila* embryos. The experimental data were digitized from Fig.3B in (51). Their best fits to Eq. 1 are plotted with solid lines. **(d)** Distance distributions between five pairs of FISH probes on chr1 in fibroblast cells. The experimental data (histograms) were digitized from Fig.4B in (52). The fits using Eq. 1 are plotted with solid lines. **(e)** Distance distributions between seven pairs of FISH probes in the Tsix/Xist region on the X chromosome of mouse ESC. The experimental data (black lines) were digitized from Fig.2F in (53), and their corresponding fits are shown in red.

**Fig. S2.**
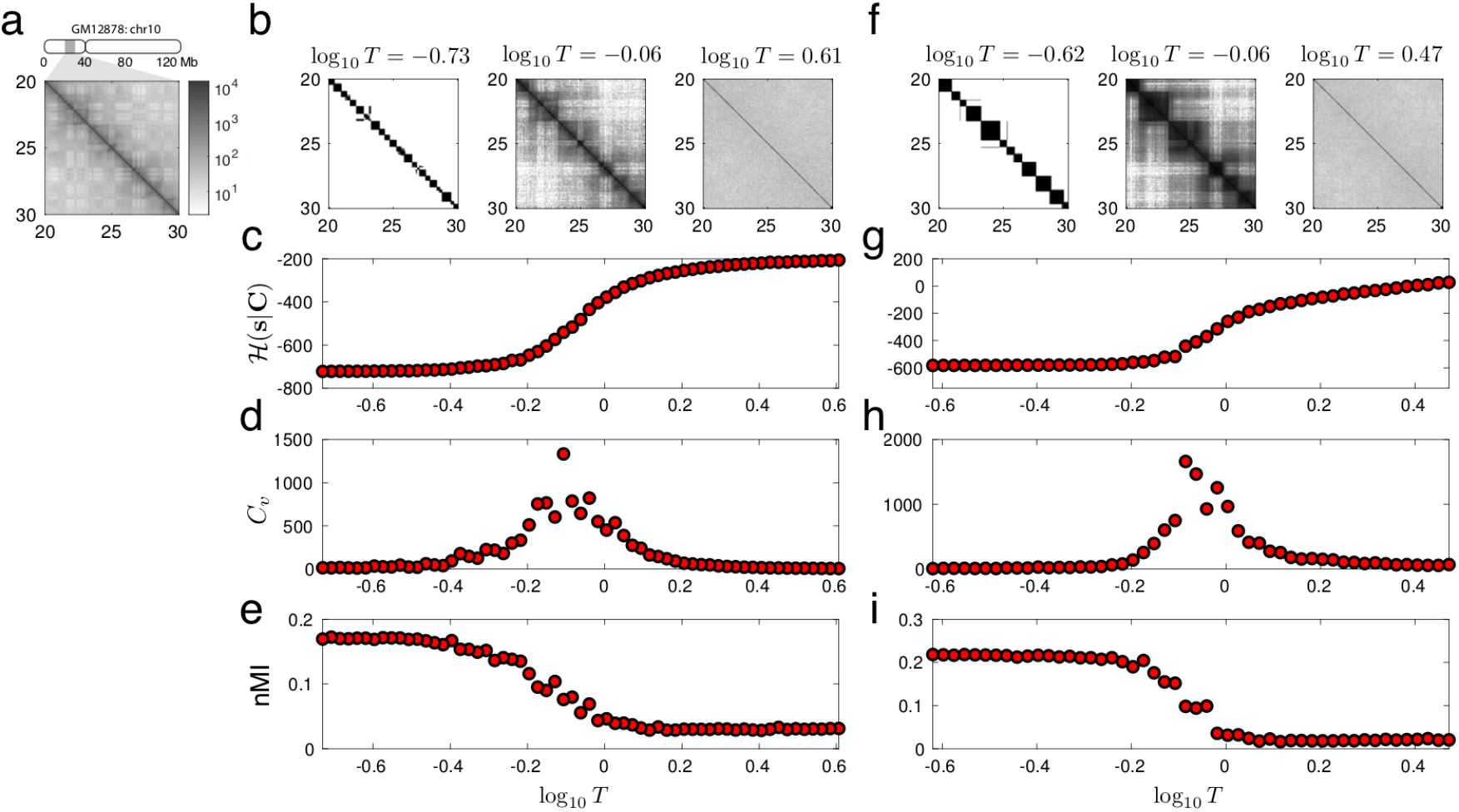
Finding the best domain solution through simulated annealing. **(a)** A subset of Hi-C data, covering 10-Mb genomic region on chr10 of GM12878. **(b)** CD solutions, obtained from the Hi-C data in **(a)**, at three values of *T* for *λ* = 0. The CD solution at each *T* was constructed by 2, 000 sample trajectories being equilibrated. **(c-e)** We plot three quantities over varying *T*, where the simulated annealing from high to low *T* (right to left in figure) was used as a sampling protocol. **(c)** The effective energy hamiltonian 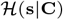. **(d)** The heat capacity 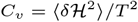. **(e)** The normalized mutual information (nMI) between the domain solution and Hi-C matrix (log_10_ **M**). **(f-i)** Same analyses repeated for *λ* = 10.

**Fig. S3.**
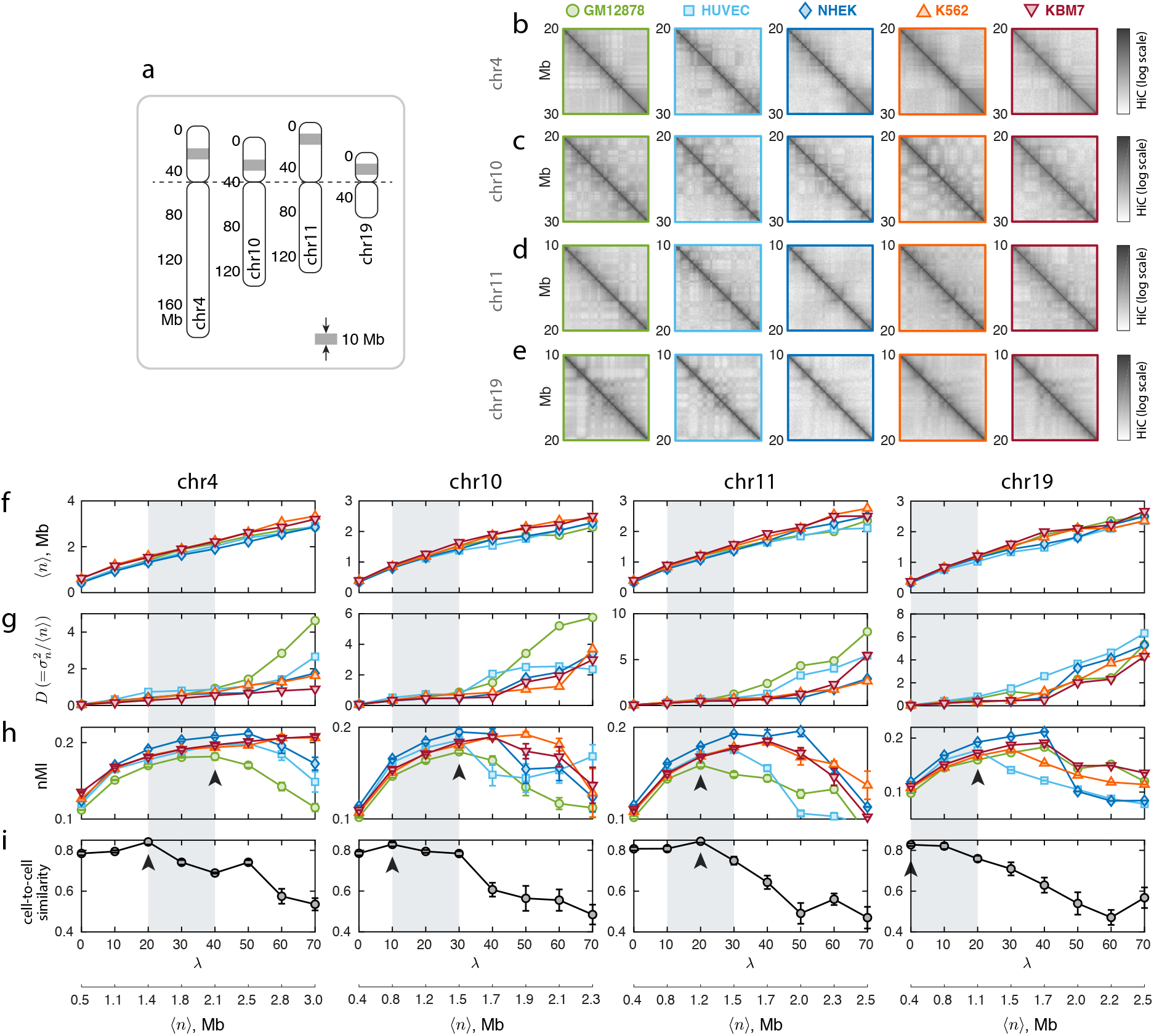
Chromatin domain solutions for chromosomes 4, 10, 11 and 19. **(a)** Relative sizes of chromosomes considered, aligned at the centromeres. The gray shade in each chromosome indicates the 10-Mb interval for which we show the Hi-C data in the next panels. **(b-e)** Hi-C data for the corresponding 10-Mb genomic intervals of **(b)** chr4, **(c)** chr10, **(d)** chr11, and **(e)** chr19, for the five different cell lines respectively. All the panels for chr10 are reprints of Fig. 3 in the main text. **(f-i)** Statistics of the domain solutions for chr4, chr10, chr11, and chr19. The five cell lines are color coded as indicated at the top of **(b)**. **(f)** Mean domain size ⟨*n*⟩ as a function of *λ*. **(g)** The index of dispersion 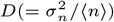 of domain sizes. **(h)** The goodness of domain solutions, measured in terms of the normalized mutual information with respect to Hi-C data (log_10_ **M**). **(i)** The similarity of domain solutions across the five different cell types, measured by the Pearson correlation between binarized contact matrices. For each chromosome, arrows indicate the likely TAD scale (highest cell-to-cell similarity) and the likely meta-TAD scale (where the nMI is high and the index of dispersion *D* starts to diverge).

**Fig. S4.**
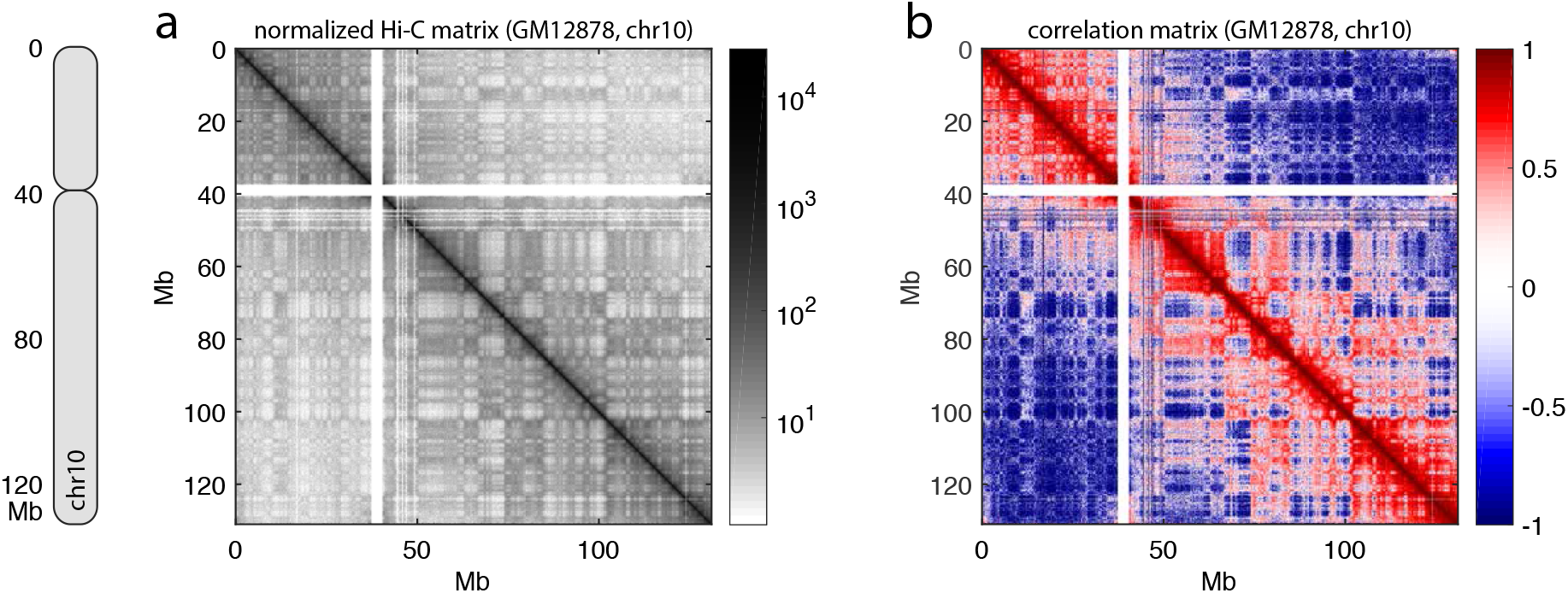
A full-chromosome view of the input data, for the chromosome 10 of GM12878. **(a)** the normalized Hi-C matrix **M**, and **(b)** the correlation matrix **C**.

**Fig. S5.**
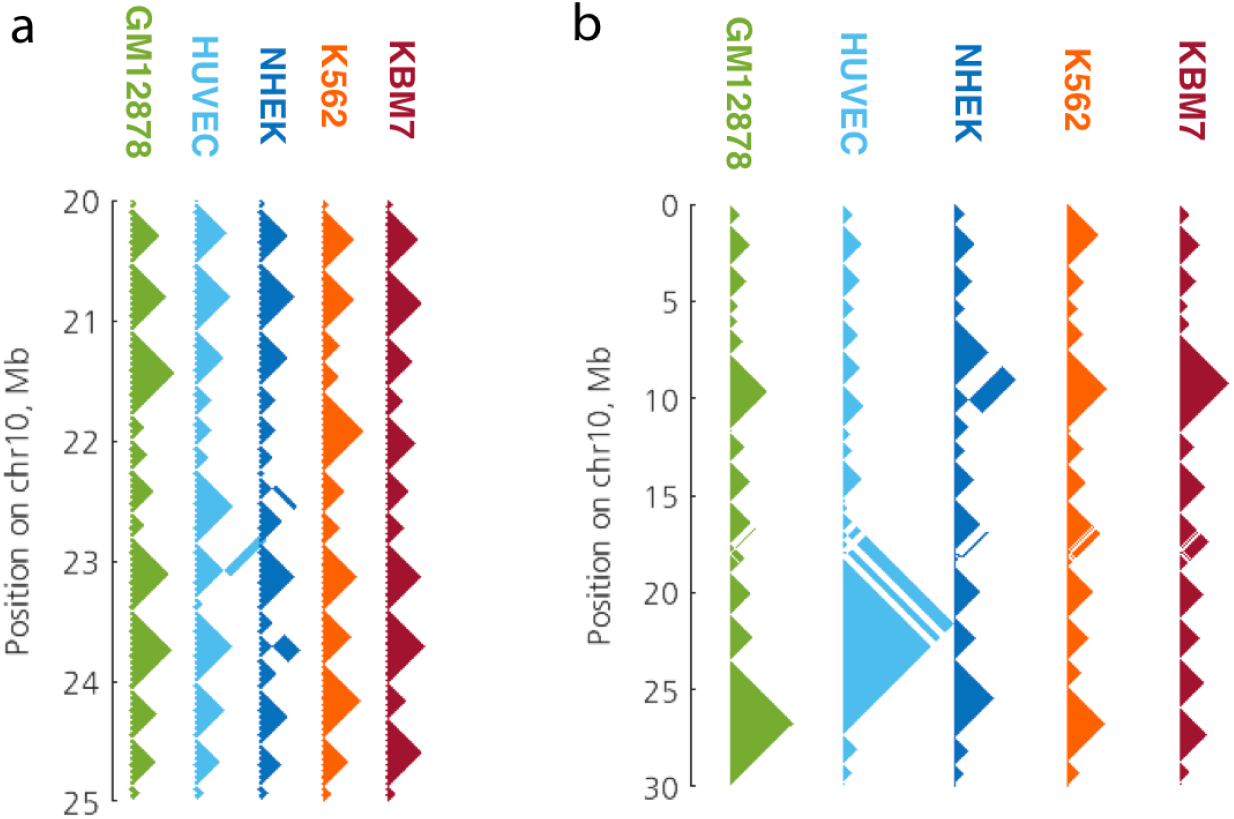
Examples of Multi-CD domain solutions at different scales. Shown are the domain solutions obtained from Multi-CD for the five different cell lines (GM12878, HUVEC, NHEK, K562, KBM7), at (a) *λ* = 0 and (b) *λ* = 40.

**Fig. S6.**
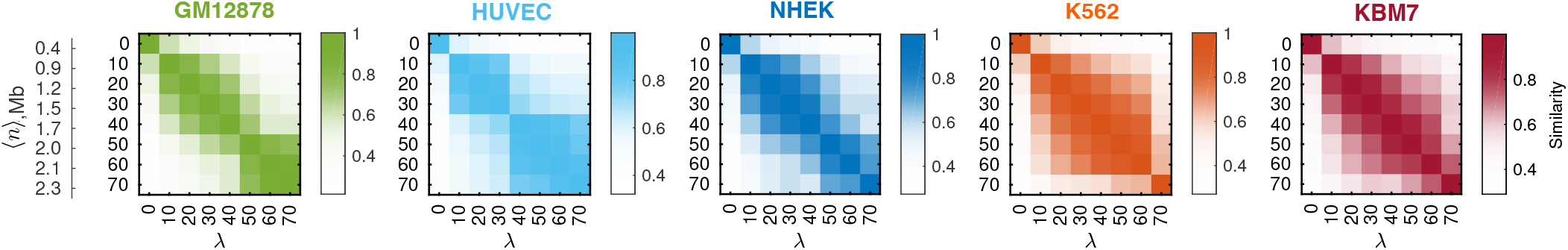
Scale-to-scale similarity of domain solutions for different cell lines. We calculate the similarity between domain solutions at different *λ* in terms of Pearson correlation. The calculation was performed for chromosome 10 from five different cell lines.

**Fig. S7.**
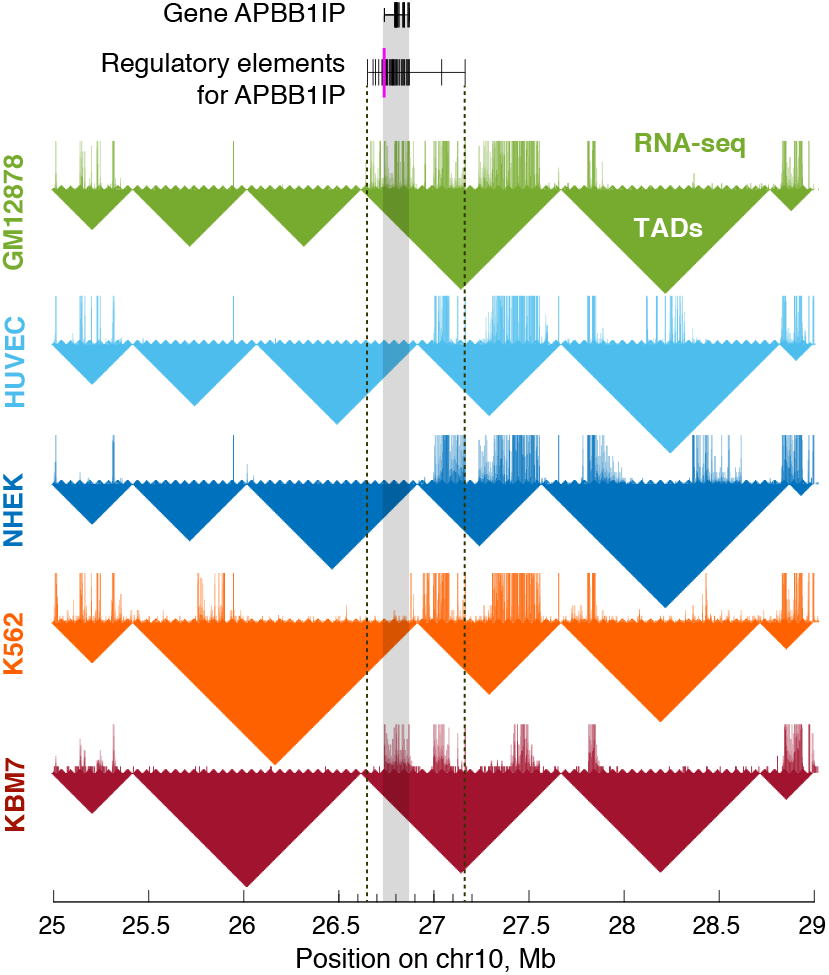
Cell-line dependent TAD organization and its link to gene expression. The RNA-seq signals from five different cell lines (colored hairy lines) are shown on top of the TAD solutions obtained by Multi-CD (triangles with matching colors). At the top shown are the the position of a specific gene APBB1IP (top row), and the regulatory elements associated with this gene (second row), including the enhancers and the promoter (the position of promotor is marked with a magenta line). APBB1IP is transcriptionally active only in two cell lines, GM12878 and KBM7. In the two cell lines, the regulatory elements are fully enclosed in the same TAD.

**Fig. S8.**
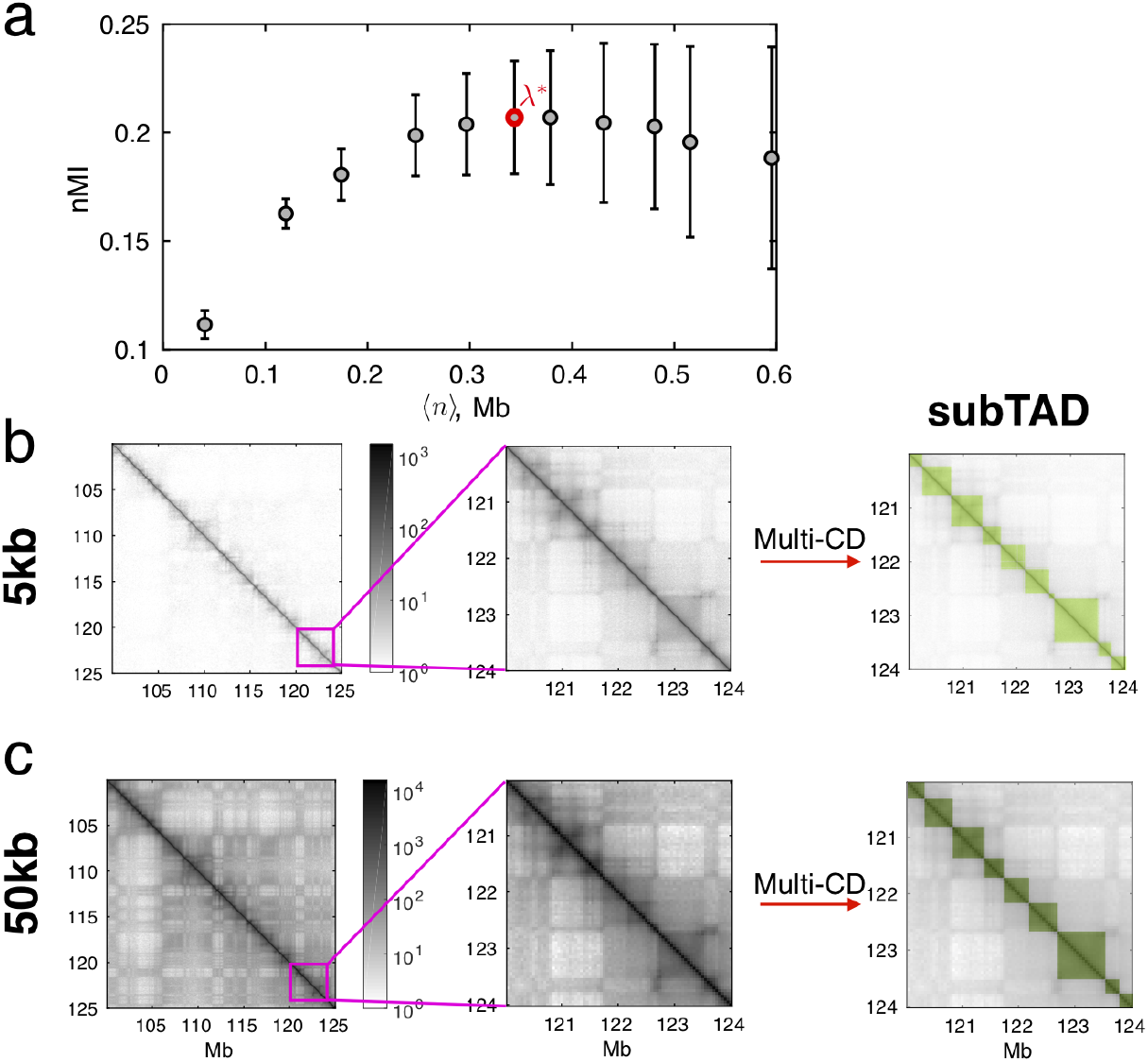
Identification of sub-TAD boundaries at 5-kb resolution. **(a)** The optimum cluster size, best describing 5-kb resolution Hi-C map in terms of nMI, is determined at ⟨*n*⟩ = 0.35 Mb, which is consistent with the sub-TAD size determined from 50-kb resolution Hi-C at *λ* = 0. **(b-c)** Comparison between Multi-CD solutions at different resolutions of the input Hi-C data, that point to the robustness of sub-TAD boundaries regardless of Hi-C resolution. **(b)** The best CD solution (corresponding to *λ* = *λ*\* in panel **(a)**) for the 5-kb resolution Hi-C data in the 120-124 Mb region of the genome. **(c)** Solution for the same genomic interval from 50-kb Hi-C, determined at *λ* = 0. The two CD solutions are effectively identical, which supports our interpretation of sub-TAD as the unit of hierarchical chromosome organization.

**Fig. S9.**
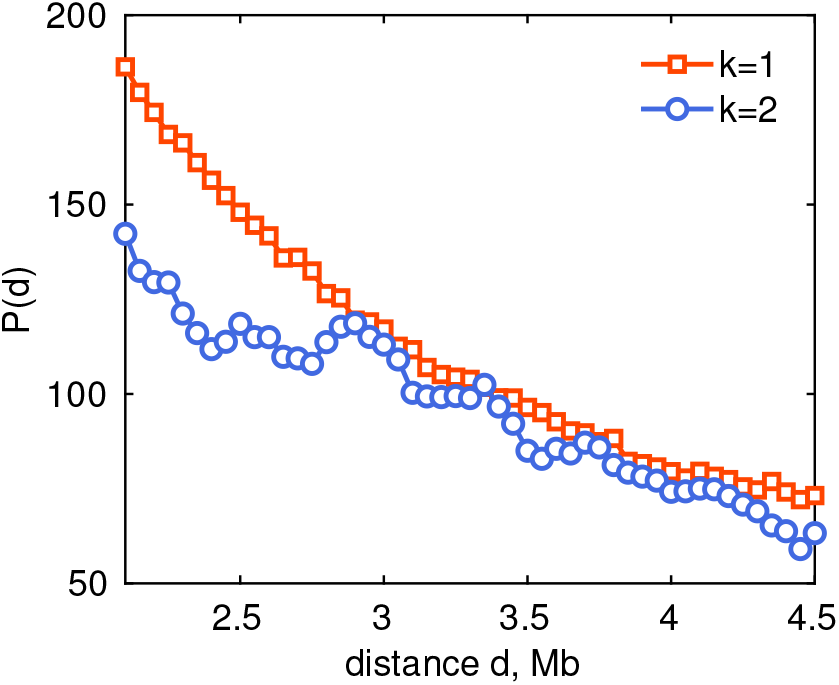
Intra-domain contact profiles for the two compartment solutions. We plot the genomic distance-dependent contact number for the two large domains *k* = 1 and *k* = 2, from the solution at *λ* = 90 for chromosome 10 in GM12878. At short genomic distance, the domain solution of *k* = 1 is characterized with a greater number of contacts than *k* = 2, which suggests that *k* = 1 domain is locally more compact. We therefore associate the first domain *k* = 1 to the B-compartment, and the second domain *k* = 2 to the A-compartment.

**Fig. S10.**
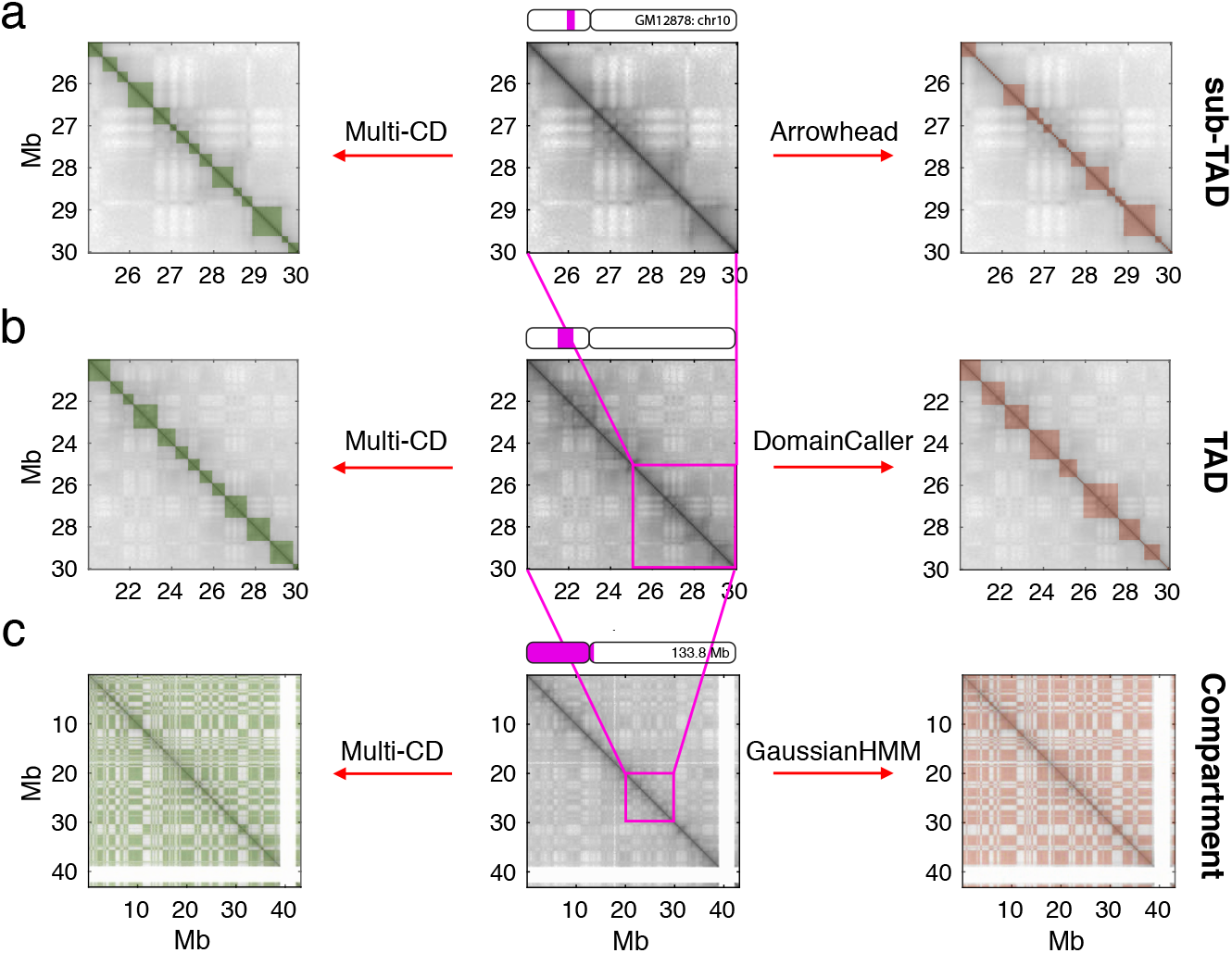
Comparison of domain solutions from Multi-CD and other methods at specific scales. Comparison between domain solutions obtained by three popular algorithms (ArrowHead, DomainCaller, GaussianHMM) (right column) and those by Multi-CD (left column), applied to 50-kb resolution Hi-C data. Three subsets from the same Hi-C data (log_10_ **M**), with different magnification (5, 10, and 40 Mb from top to bottom), are given in the middle column. ArrowHead algorithm (19) was used for identifying the domain structures of sub-TADs, DomainCaller (22) for TADs, and Gaussian Hidden Markov Model (GaussianHMM) (19) for compartments. Multi-CD use *λ* = 0, 10, 90, as the parameter values for identifying sub-TADs, TADs, and compartments, respectively.

**Fig. S11.**
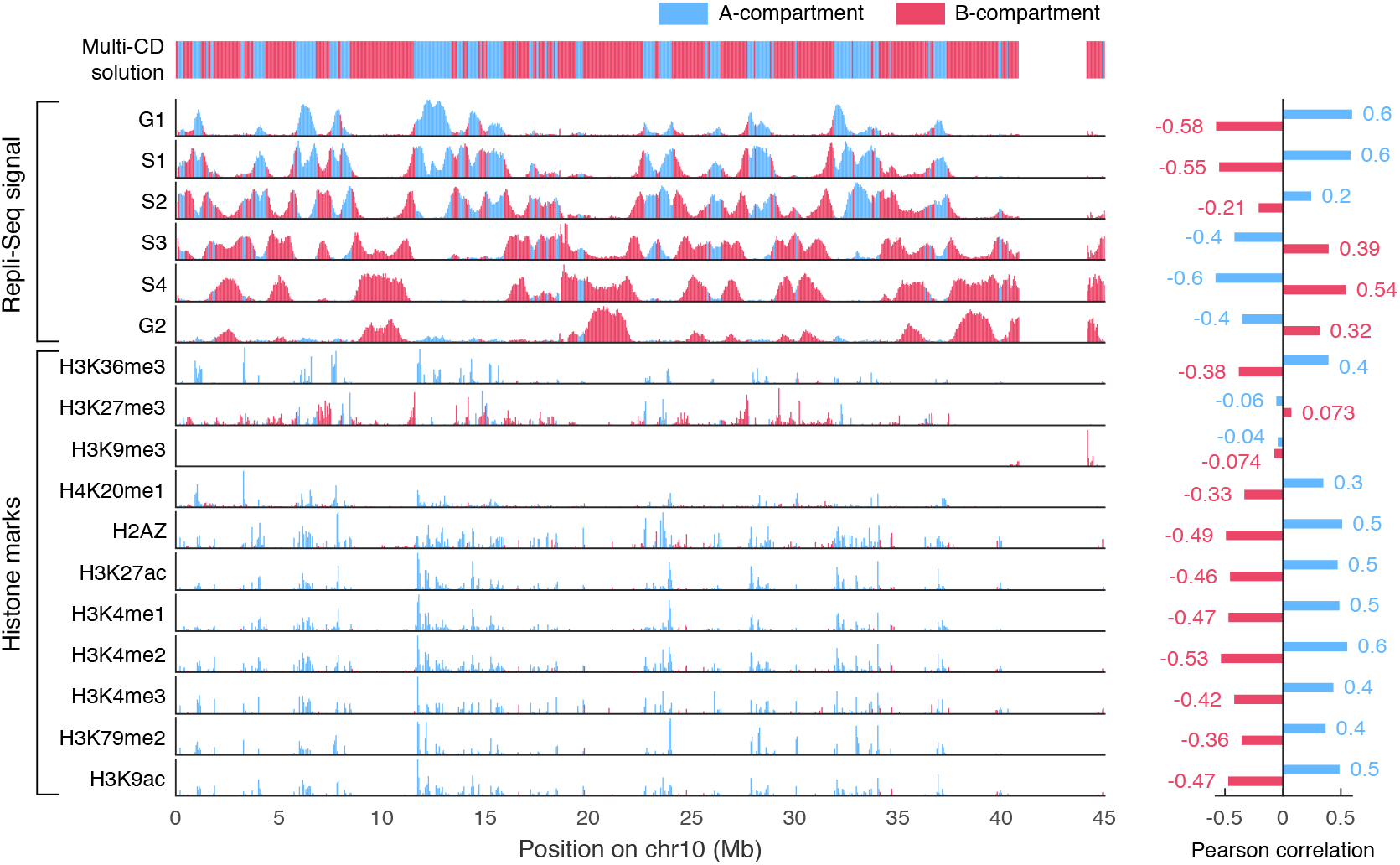
Comparison of histone marks and compartments. Extension of Fig. 6c-d which make comparison between the CD solutions for A/B-compartments by Multi-CD and epigenetic marks. The upper part with Repli-Seq signals is a reprint from the main text figure. The lower part shows histone marks on the corresponding genomic range. Majority of the histone marks are correlated with the A-compartment. The values of Pearson correlation between Repli-Seq signal or histone marks and A/B-compartment are given on the right.

**Fig. S12.**
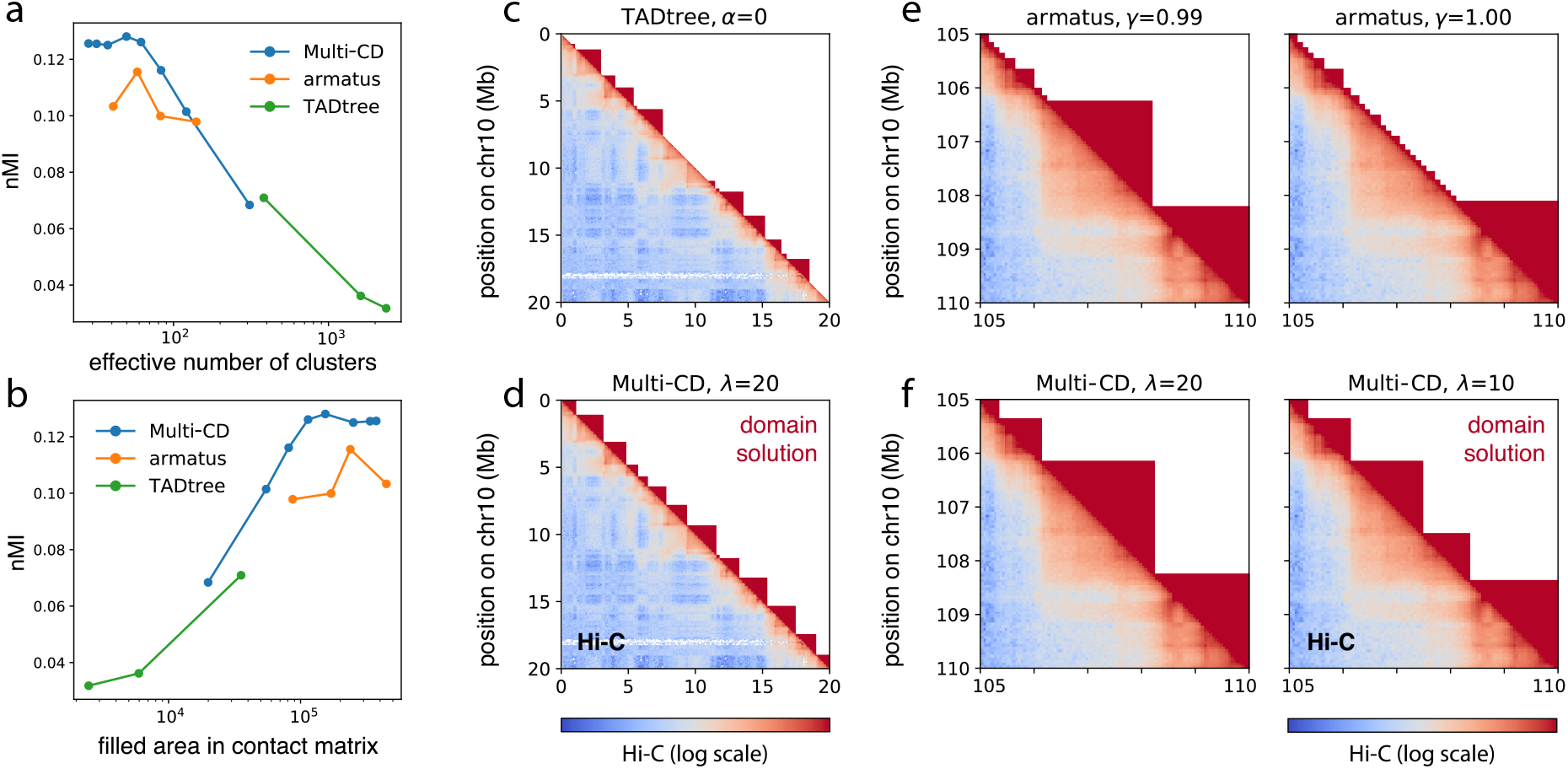
Comparison to existing algorithms for identifying domains at multiple scales. **(a,b)** Normalized mutual information between domain solutions at multiple scales, from Multi-CD, Armatus (33) and TADtree (32) respectively, and the log10 of KR-normalized Hi-C matrix for chr10 of the cell line GM12878. The scale of a domain solution **s** is measured in two ways, in terms of **(a)** the effective number of clusters, 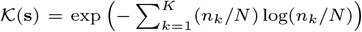, where 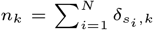 is the domain size; and **(b)** the total area of 1’s in the corresponding binary contact matrix, 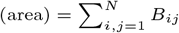 where 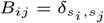. All domain solutions from TADtree and Armatus were obtained using the respective default parameter settings. **(c-f)** Visual comparison of domains found by **(c,d)** TADtree and Multi-CD, and **(e,f)** Armatus and Multi-CD, at matching scales in terms of the average domain size. Domain solutions are shown in the upper triangle, colored by red (intra-domain) and white (extra-domain) for effective visualization. The lower triangle plots the corresponding subset of the Hi-C data (KR-normalized and in log10). Refer to the original papers (32, 33) for the definitions of the respective control parameters *α* (TADtree) and *γ* (Armatus).

**Fig. S13.**
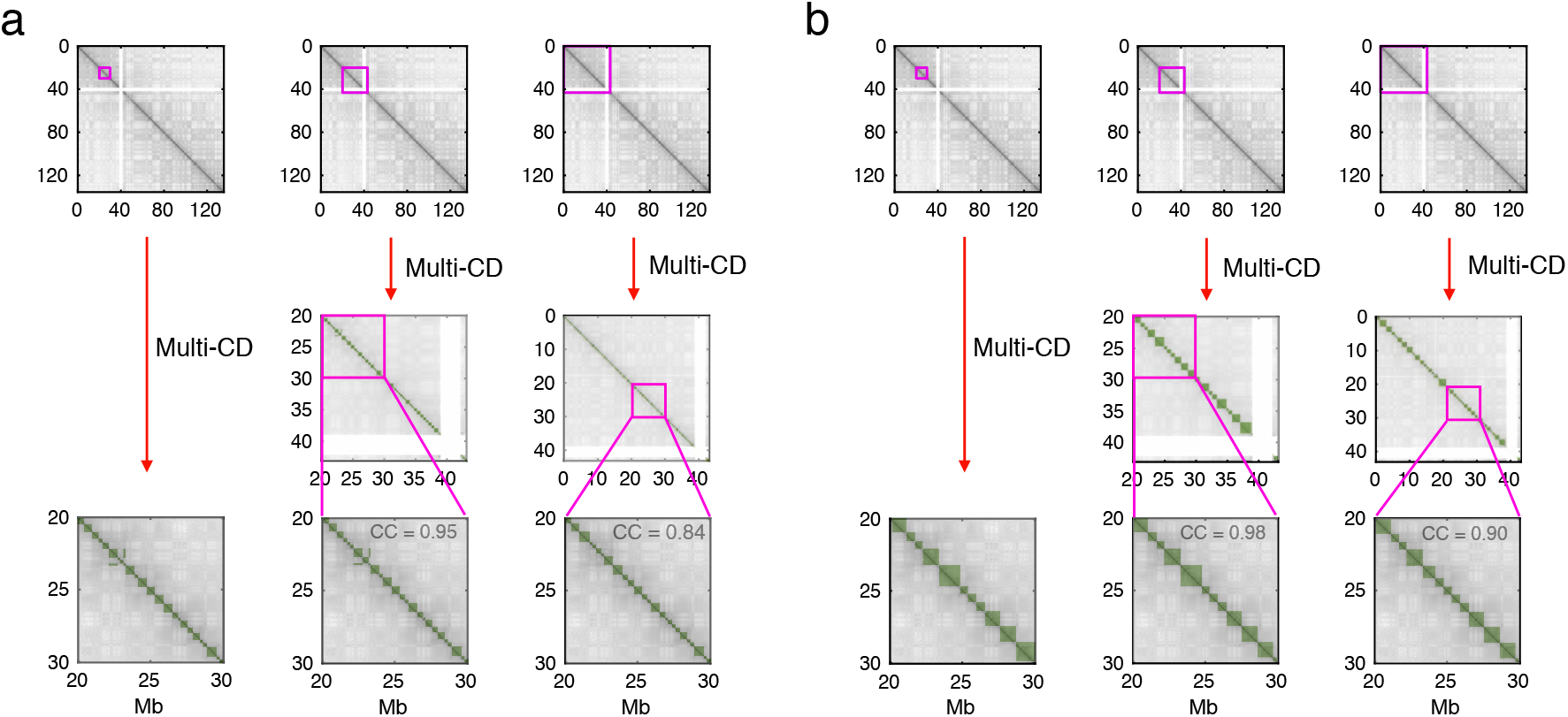
Robustness of clustering solutions over different subsets of Hi-C data. Here we compare domain solutions from Hi-C inputs of different size. Multi-CD is confirmed to be *locality-preserving*. That is, the sets of domain solutions determined from Hi-C inputs with different sizes remain almost identical to each other. The Hi-C data demarcated by the purple squares on the top panels are the input data used for Multi-CD analysis. The three panels from left to right on the bottom are the domain solutions from 10-Mb, 20-Mb, and 40-Mb Hi-C inputs. **(a)** For *λ* = 0, the correlation coefficients of 20-Mb Hi-C and 40-Mb Hi-C generated domain solutions with respect to the 10-Mb Hi-C generated one is 0.95 and 0.84, respectively. **(b)** Same calculations were carried out for *λ*=10.

**Fig. S14.**
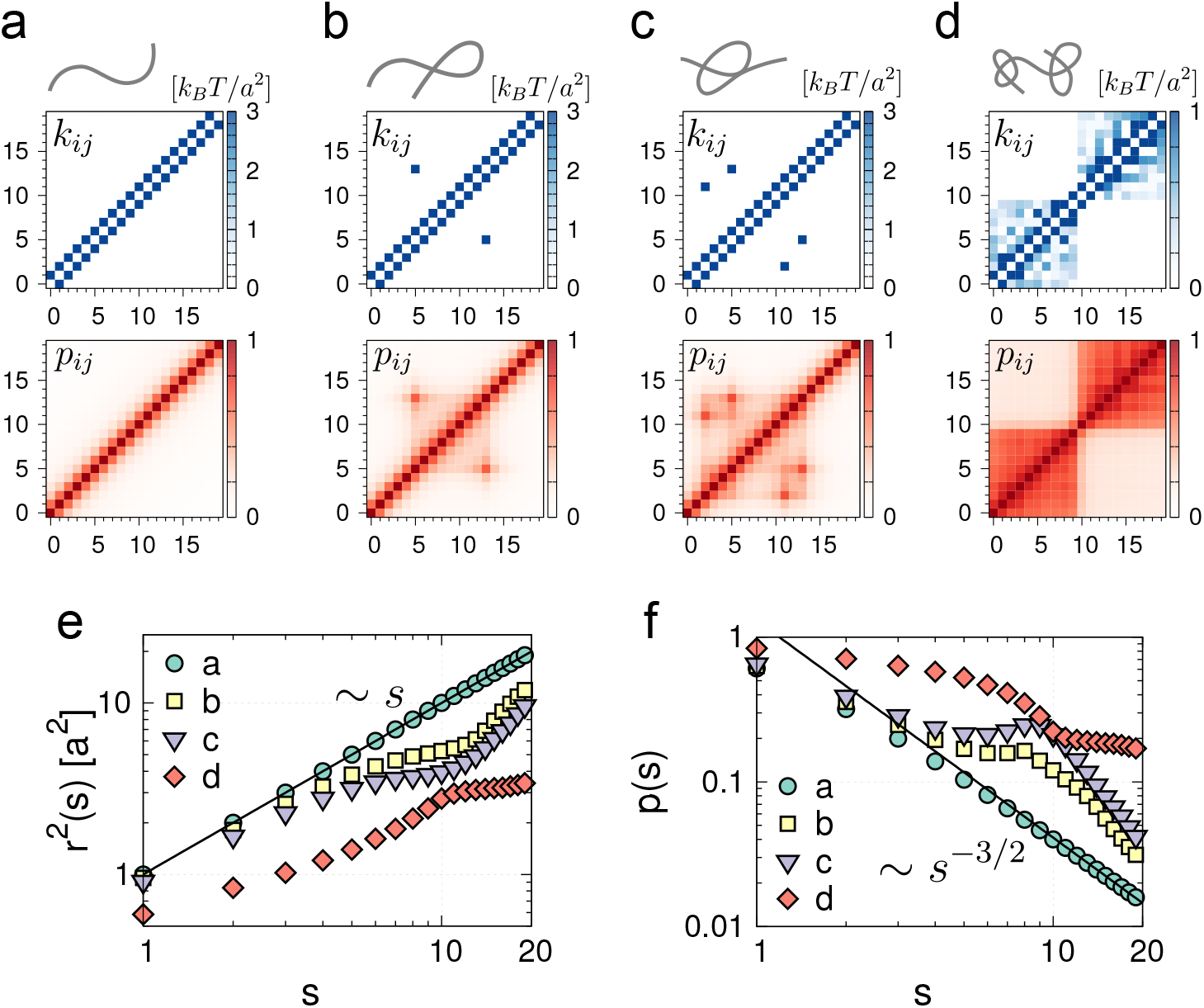
Heterogenous loop model (14) to compare the contact probabilities of Gaussian polymer networks. **(a-d)** Four examples of polymer models composed of 20 monomers with different interaction strength matrix [*k_ij_*] (top row), and the corresponding contact probability matrices [*p_ij_*] (second row) calculated with *r_c_* = 1. **(e)** the mean square distance and **(f)** the contact probability *p*(*s*) are calculated as a function of the genomic distance, s, for the four different models (a-d). Scaling results in (e) and (f) show that even the Gaussian polymer network model can produce rich multi-scale structure with domains.

